# RBM39 degrader invigorates natural killer cells to eradicate neuroblastoma despite cancer cell plasticity

**DOI:** 10.1101/2024.03.21.586157

**Authors:** Shivendra Singh, Jie Fang, Hongjian Jin, Lee-Ann Van de Velde, Qiong Wu, Andrew Cortes, Christopher L. Morton, Mary A. Woolard, Waise Quarni, Jacob A. Steele, Jon P. Connelly, Liusheng He, Rebecca Thorne, Gregory Turner, Thomas Confer, Melissa Johnson, William V. Caufield, Burgess B. Freeman, Timothy Lockey, Shondra M. Pruett-Miller, Ruoning Wang, Andrew M. Davidoff, Paul G. Thomas, Jun Yang

**Author notes:** These authors contributed equally (Shivendra Singh, Jie Fang, Hongjian Jin, Lee-Ann Van de Velde).

## Abstract

The cellular plasticity of neuroblastoma is defined by a mixture of two major cell states, adrenergic (ADRN) and mesenchymal (MES), which may contribute to therapy resistance. However, how neuroblastoma cells switch cellular states during therapy remains largely unknown and how to eradicate neuroblastoma regardless of their cell states is a clinical challenge. To better understand the lineage switch of neuroblastoma in chemoresistance, we comprehensively defined the transcriptomic and epigenetic map of ADRN and MES types of neuroblastomas using human and murine models treated with indisulam, a selective RBM39 degrader. We showed that cancer cells not only undergo a bidirectional switch between ADRN and MES states, but also acquire additional cellular states, reminiscent of the developmental pliancy of neural crest cells. The lineage alterations are coupled with epigenetic reprogramming and dependency switch of lineage–specific transcription factors, epigenetic modifiers and targetable kinases. Through targeting RNA splicing, indisulam induces an inflammatory tumor microenvironment and enhances anticancer activity of natural killer cells. The combination of indisulam with anti-GD2 immunotherapy results in a durable, complete response in high-risk transgenic neuroblastoma models, providing an innovative, rational therapeutic approach to eradicate tumor cells regardless of their potential to switch cell states.

## INTRODUCTION

Pediatric cancers give rise to unique challenges in the clinic. First, the number of potentially “druggable” targets that can be acted upon with specific and selective inhibitors remains low^1,2^. Second, the alterations of cancer signaling pathways have not proven to be viable therapeutic targets ^3–6^. Third, the low mutation burden of pediatric tumors leads to a low spectrum of neoantigens^7^, which may greatly limit the effect of immunotherapy including immune checkpoint inhibitors. Molecular and cellular heterogeneity within a tumor causes further challenges for designing and selecting effective therapies, and curtailing treatment resistance^8^. Using neuroblastoma as an example, this study provides evidence that heterogenous tumor cells can be eliminated by harnessing the power of the immune system through targeting the RNA splicing factor, RBM39.

Arising in early fetal development^9^, neuroblastoma is an extracranial, embryonal tumor derived from the neural crest lineage, transformed by *MYC* oncogenes^10–13^. 50% of high-risk neuroblastomas have *MYCN* amplification, while the other half express high levels of *C-MYC*. With current intensive multimodal therapies (combined cytotoxic chemotherapies, surgery, stem cell transplantation, radiotherapy, differentiating agents and anti-GD2–based immunotherapy), 5-year survival rates for patients with high-risk neuroblastoma remain less than 50%^14–17^. In addition, survivors of high-risk disease have a significant risk of developing long-term side effects, including subsequent malignant neoplasms due to exposure to cytotoxic chemotherapy and radiotherapy^18,19^. Unfortunately, developing safer and more effective targeted therapies against high-risk neuroblastoma has been challenging due to the low incidence of targetable recurrent mutations^20–22^, although a small fraction of patients with ALK mutations are highly responsive to ALK inhibitors^23^. While functional genomic screens have identified many essential survival genes in neuroblastoma cells^24–29^, translating these discoveries into effective therapies has been technically challenging because most of these essential genes, including MYC, are considered to be “undruggable”. To address the unmet clinical need for high-risk neuroblastoma therapy, we recently identified a new therapeutic target, RBM39, a MYC target that regulates pre-mRNA splicing and appears to be essential to neuroblastoma survival^30^. We and others further found that indisulam, a “molecular glue” drug that selectively recruits RBM39 to the CRL4-DCAF15 E3 ubiquitin ligase (DCAF15) for proteasomal degradation^31,32^, is highly effective against neuroblastoma by inducing a wide range of splicing anomalies that affect a number of essential genes^30,33^. It is known that MYC– driven neuroblastoma has deregulated splicing^34–36^. However, high-risk, patient-derived xenograft (PDX) models implanted in immune deficient mice eventually relapse when treated with this RMB39 degrader, prompting us to investigate the mechanism of therapy resistance of neuroblastoma to indisulam treatment.

Lineage plasticity^37^ (or transdifferentiation^38^, pathway indifference^39^) is one key mechanism of drug resistance by which cancer cells acquire an alternative cellular state in response to treatment to sustain their survival^37–41^. Neuroblastoma cell lines contain morphological variants that contribute to cell state heterogeneity^42–44^, with cells residing in one of two major cellular states, committed adrenergic (ADRN) and neural crest migratory or mesenchymal (MES), largely defined by epigenetic and transcriptomic programs^45–47^. These distinct populations exhibit differential responses to chemotherapy^46^. While there is a theory that MES and ADRN states are interconvertible, consequently leading to therapy resistance, this is largely supported by forced overexpression or genetic deletion of transcriptional factors in cell lines^46,48,49^. Whether interconversion of these cell states actually occurs in response to tumor environmental changes such as exposure to therapeutic agents is still not very clear, albeit these studies indicate that tumor cells acquired MES gene features when developed resistance. Neuroblastoma results from differentiation arrest of neural crest−derived sympathoadrenal progenitor cells^10,50^. Recent single cell RNA-seq and lineage tracing studies show transcriptomic similarity between neuroblastoma and normal cell types along the developmental trajectory of neural crest cells^51–54^. The differentiation status of human adrenal medulla at different developmental time points is associated with neuroblastoma outcome risk^55^. Unlike cell lines which can be clearly defined by ADRN and MES signatures, the *in vivo* cellular state of neuroblastoma cells is less clear, although most resemble an ADRN state^51,54,56^. However, malignant cells of high-risk neuroblastomas show increased MES signatures and reduced ADRN signatures in one study^52^. Other studies further showed that ADRN tumors with MES features have molecular traits of Schwann cell precursors (SCP), bridge cells and early neuroblasts (all of which are neural crest cell progeny)^51,52^, indicating intratumoral plasticity in those tumors. However, the path from one cellular state of neuroblastoma to another (or others) during *in vivo* therapy has not been defined. Our study addresses one key knowledge gap in therapy resistance: how cellular plasticity of cancer cells confers drug resistance in neuroblastoma. Addressing this question is a key step towards developing more effective combination therapies against refractory malignancies.

Here, by using multiple high-risk neuroblastoma models that developed resistance to indisulam, we show that cancer cells undergo a multi-directional switch of cell states, reminiscent of the cellular pliancy of neural crest cells. The lineage switch of neuroblastoma cells is associated with epigenetic changes, and dependency switch of lineage–specific transcription factors, epigenetic modifiers and targetable kinases. Notably, indisulam treatment induces an inflammatory tumor microenvironment characterized by infiltration of immune cells such as T and NK cells. In the NK cell competent mice that are deficient in adaptive immunity (T cells and B cells), implanted c-MYC–driven murine neuroblastomas are completely eradicated by indisulam, suggesting that NK cells may play a critical role in indisulam-mediated anticancer activity. Complete durable responses are also achieved in an immune competent MYCN/ALK^F1178L^ neuroblastoma mouse model when treated with the combination of indisulam and anti-GD2 mAb. It is known that anti-GD2 mAb exerts its antitumor effect through NK cell-mediated ADCC (antibody dependent cell–mediated cytotoxicity)^57^. Our study indicates that targeting RBM39 in combination with anti-GD2 therapy may eradicate neuroblastoma cells irrespective of their cell state switching potential, providing a rationale to combine indisulam and anti-GD2 mAb as a new therapy for high-risk neuroblastoma patients.

## RESULTS

### Indisulam resistance in a transgenic *MYCN/ALK^F^*^1178^*^L^*mouse model is associated with RBM39 upregulation and a lineage switch to MES and Schwann cell precursor phenotypes

Human *ALK^F1174L^* (analogous to mouse *ALK^F1178L^*) is the most frequent somatic mutation in neuroblastoma and is associated with *MYCN* amplification, conferring a worse prognosis than *MYCN* amplification alone^58^. Our previous study showed that indisulam treatment of *MYCN/ALK^F1178L^* mice with a regular dosing (25 mg/kg, 5 days on/2 days off for two weeks) led to durable, complete responses even when tumor sizes reached over 1000mm^3^ prior to treatment initiation^30^. When testing the dosing schedule by treating the *MYCN/ALK^F1178L^* mice with only one dose of indisulam (25 mg/kg) per week for 3 weeks in our current study, we found variable responses (**Figure 1A**). Two weeks after therapy discontinuation, we resumed the regular dosing schedule for an additional two weeks. Among the 5 treated mice, three showed complete and durable responses, one responded to the first dose of therapy (tumor volume reduced from 1100mm^3^ to 300mm^3^) but then developed resistance to the following treatment (**Figure 1A**), while another one showed a complete response but eventually relapsed after second therapy discontinuation (**Figures 1A and 1B**). To understand the mechanism of therapy resistance, we performed RNA-seq analysis followed by gene set enrichment analysis (GSEA) to compare the naïve tumors with the relapsed and resistant tumors (**Figure 1C**). In comparison with the naïve tumors that showed a high “E2F” gene signature, indicative of the high proliferation rate of naïve tumor cells, the “human 20q11 amplicon” was the top upregulated gene signature in the resistant/relapsed tumors (**Figures 1C and 1D**). Human *RBM39* is located in 20q11, suggesting that there was a selective pressure under indisulam treatment leading to high levels of expression of *Rbm39* locus in mouse. This hypothesis was consistent with the high levels of RNA-seq reads of *Rbm39* in the relapsed and resistant tumors, which is located in Chr2qC of mouse genome (**Figure 1E**). This was further verified by the correlation of indisulam resistance with the RBM39 copy number and mRNA expression levels across 534 cell lines in DepMap (depmap.org) (**Figures 1F and 1G**).

**Figure 1.**
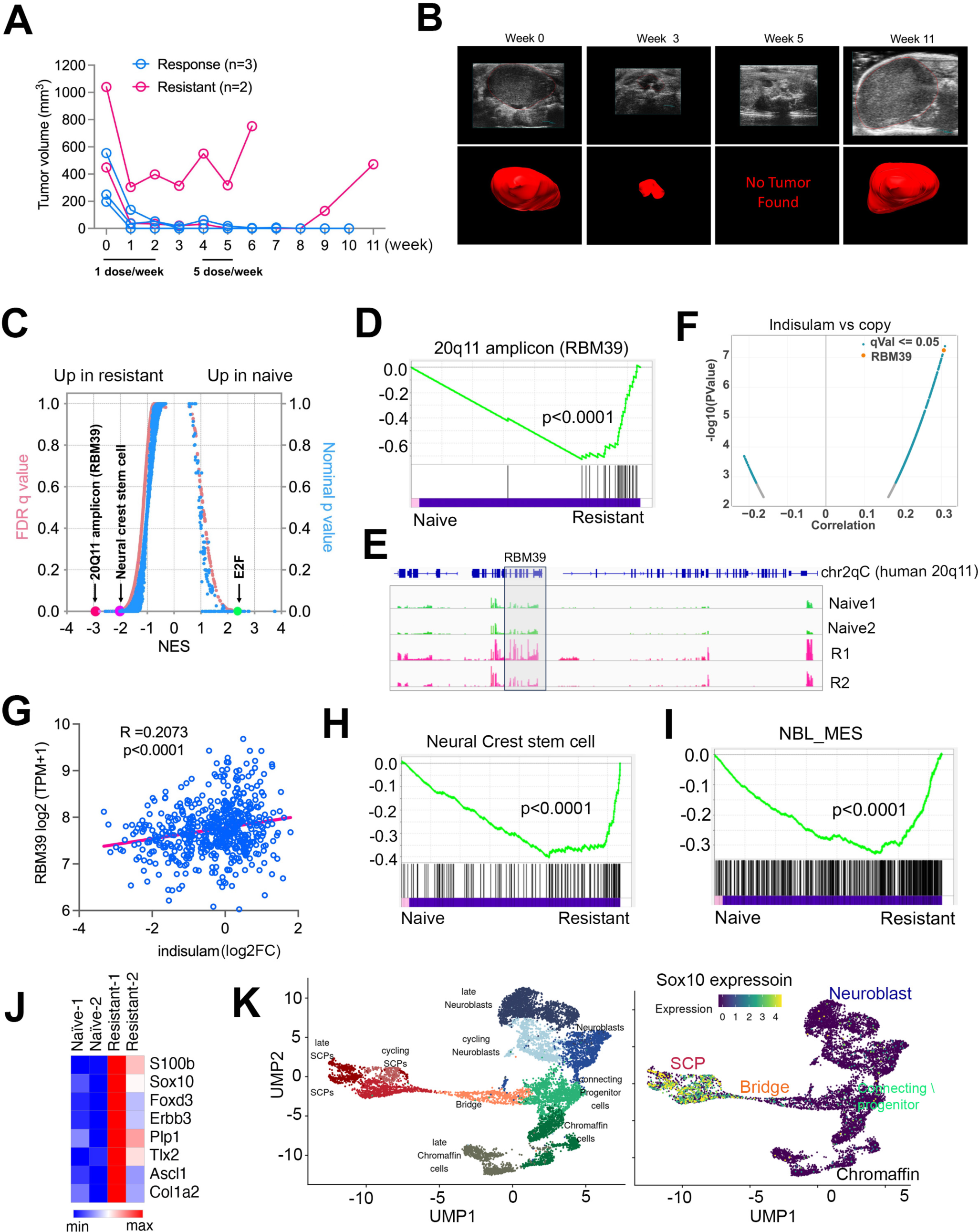
Indisulam resistance in transgenic *MYCN/ALK^F1178L^* mouse models is associated with *Rbm39* upregulation, and lineage switch to cell states with MES and Schwan precursor cells. (A) Tumor growth curve for five *MYCN/ALK^F1178L^* mice treated with 25 mg/kg of indisulam. Blue color indicates sustained complete response (n=3). Pink color indicates tumor resistance and relapse (n=2) (B) Ultrasound imaging demonstrates one example of tumor relapse. Top panel shows the two-dimensional ultrasound images and the bottom panel shows the volume reconstructions. (C) Plot shows GSEA analysis results of RNA-seq data of naïve tumors vs relapsed tumors. Blue color indicates nominal P values and pink color indicates false discovery rate (FDR) q values. (D) GSEA plot shows “20q11 amplicon” gene set is significantly upregulated in resistant tumors. (E) Plot shows the normalized RNA-seq reads at *Rbm39* gene locus by IGV program. (F) Plot shows the Pearson correlation between gene copy number vs indisulam response from DepMAP data. (G) Plot shows the Pearson correlation between gene expression vs indisulam response from DepMAP data. (H, I) GSEA plots shows “neural stem cell” and “NBL_MES” gene sets are significantly upregulated in resistant tumors. (J) Heatmap shows the expression of Schwan cell progenitor (SCP) markers in naïve tumors vs relapsed tumors. (K) *Sox10* labels the SCP cell population as the progenitors of neuroblasts and Chromaffin cells that are origin of neuroblastoma. Left shows the distinct cell populations in human adrenal medulla and the right shows the expression of *Sox10* in each cell population, based on scRNA-seq analysis from the study^9^.

Additionally, we noticed that the relapsed/resistant tumor cells acquired a high “neural crest stem cells” gene signature (**Figures 1C and 1H**), suggesting that these therapy resistant cancer cells underwent de-differentiation. We then examined the ADRN and MES signatures^46^ and found that the relapsed/resistant cells exhibited a significant shift to the MES state (**Figure 1I**). Particularly, we also found that the relapsed/resistant tumor cells expressed a signature of “Schwann cell precursors (SCP)”^51,52^, characterized by high expression of *S100b*, *Sox10*, *Plp1* and other genes (**Figure 1J**). Then we projected the expression of *Sox10*, the SCP transcription factor, into the single cell analysis of human adrenal medulla, which demonstrated high levels of *SOX10* expression in SCPs but not in chromaffin and neuroblast cells (**Figure 1K**), the downstream progenies of SCP and origin of neuroblastoma. We further examined the SCP signature, which was highly expressed in the resistant/relapsed *Th-MYCN/ALK^F1178L^* tumors, in two human neuroblastoma cohorts (Target NBL and St Jude PCGP). The SCP markers such as *TLX3*, *SOX10, S100b*, *FOXD3* appeared to be enriched in non-*MYCN* amplified tumors (**Figures S1A, B**), suggesting that the resistant/relapsed *MYCN*-driven tumors acquired gene features similar to the non-*MYCN* amplified tumors. Taken together, these data indicate that the resistant/relapsed neuroblastomas expressed higher levels of RBM39 after they acquired new cell state properties of neural crest progenies.

### Indisulam resistance is associated with a lineage switch from ADRN to MES and melanocytic state in human *MYCN*-amplified neuroblastomas

To further understand the cell state alterations in neuroblastoma resistance, we treated *MYCN*-amplified patient-derived xenografts (SJNB14) implanted subcutaneously into CB17/SCID mice with indisulam (25 mg/kg, 5 days on/2 days off for two weeks). All tumors responded to the therapy, but eventually they relapsed. However, these tumors remained responsive to indisulam treatment until the eighth cycle of therapy when tumors developed full resistance (**Figure 2A**). RNA-seq followed by GSEA analysis showed that the resistant tumors had a significant reduction of the “ADRN” signature, followed by a “hypoxia” signature (**Figures 2B and 2C**). However, the resistant tumors acquired a high “interferon alpha and beta” signature (**Figure 2D**), in line with the feature of MES neuroblastoma cells, which were reported to express high levels of interferon pathway genes^59,60^. The master transcriptional factors of MES cells, such as *c-MYC*, *PRRX1* and *NOTCH1* were highly upregulated in the resistant tumor cells, together with the SCP marker *S100B* and stem cell markers *KIT* and *SALL4* (**Figure 2E**). Wnt ligands were also highly upregulated in the resistant tumors. A recent study based on network analysis of neuroblastoma expression data proposed that Wnt signaling is a major determinant of regulatory networks that underlie mesenchymal/neural crest cell-like cell identities through PRRX1 and YAP/TAZ transcription factors^61^. Surprisingly, melanocytic markers, including *MITF*, which encodes the master transcriptional factor of melanocytes and melanomas, were also highly upregulated. It is known that SCP is a melanocyte progenitor^62^. The scRNA-seq analysis in the developmental trajectories of SCP to chromaffin cells and neuroblasts showed no expression of *MITF* (**Figure S2A**), supporting the hypothesis that neuroblastoma cells either directly transdifferentiated to a melanocyte-like state or dedifferentiated to SCPs which then differentiated to a state with melanocytic features. We further examined the expression of Wnt ligands and melanocytic gene signatures in human neuroblastoma cohorts. The Wnt ligand signature was enriched in neuroblastomas without *MYCN* amplification in both Target NBL and St Jude PCGP cohorts^21,63^ (**Figures 2F, S2B**). Interestingly, the melanocytic gene signature was enriched in Target neuroblastomas without *MYCN* amplification but not in St Jude PCGP data (**Figures 2F, S2B**), probably due to the confounding factors such as different stages, disease risk classification, and pre-treatment and post-treatment from both cohorts. It was known that Target NBL dataset was enriched with high-stage and high-risk neuroblastomas^21^. Distinct from the *MYCN/ALK^F1178L^* resistant murine tumors, the *RBM39* expression showed no significant induction in the resistant SJNB14 tumors (**Figure 2G**), and Sanger DNA sequencing showed no mutations in the indisulam binding motif (**Figure S2C**), suggesting that the lineage switch probably plays a major role in mediating the indisulam therapy resistance in this *MYCN*-amplified PDX model.

**Figure 2.**
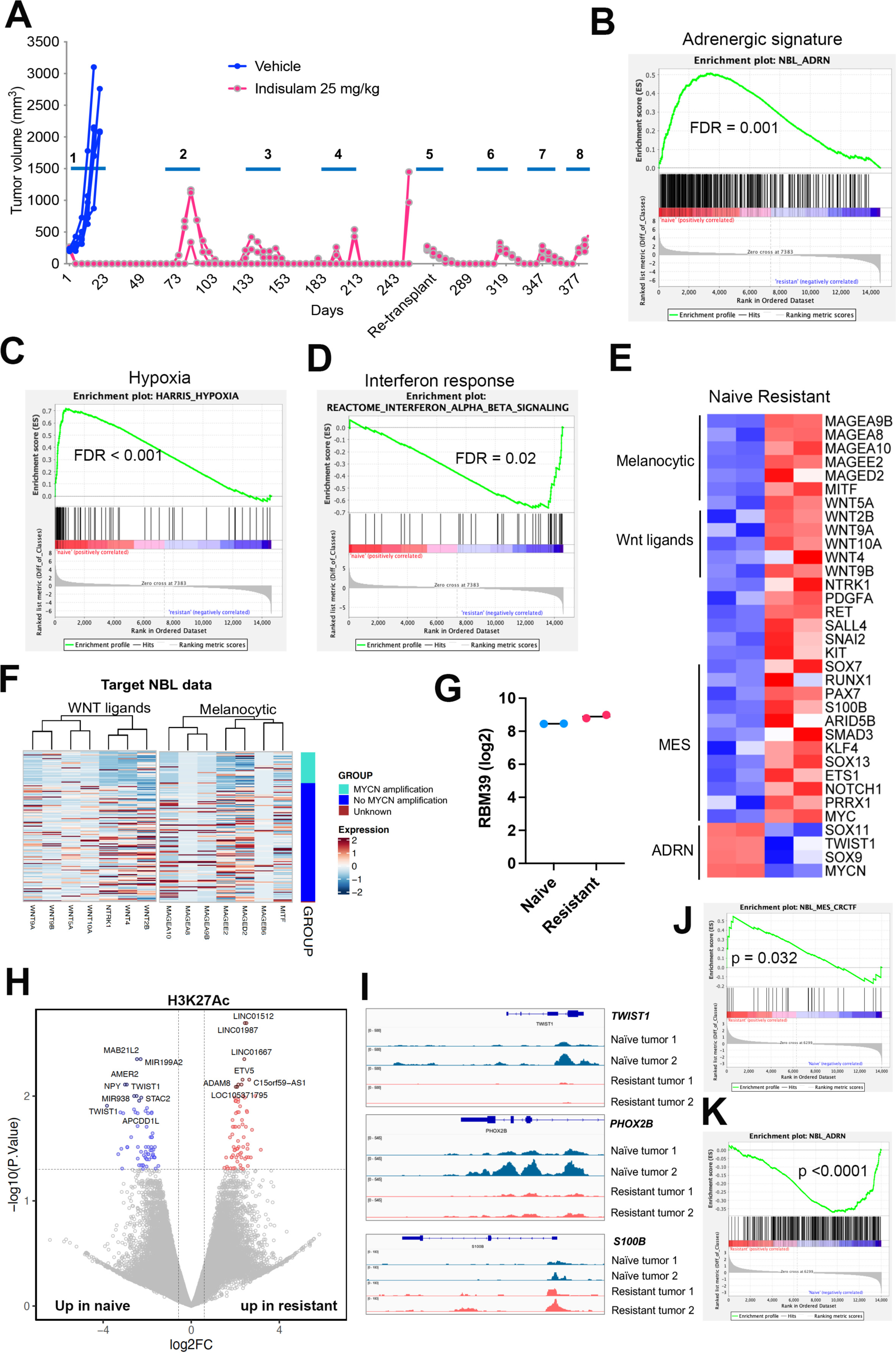
Indisulam resistance is associated with lineage switch from ADRN to MES and melanocytic state in human *MYCN*-amplified neuroblastomas. (A) SJNB14 PDX implanted in CB17/SCID mice undergoing repeated cycles (2-week as one cycle) of treatment with 25 mg/kg indisulam, 5 days on, two ways off. Note: relapsed tumors after the 4^th^ cycle were re-implanted to new mice before the 5^th^ cycle treatment due to aging of the primary mice. (B, C, D) GSEA shows the ADRN and hypoxia gene signatures are significantly downregulated in resistant tumors vs naïve tumors while the interferon gene signature is significantly upregulated. (E) Heatmap showing that the expression changes of melanocytic markers, Wnt ligands, MES and ADRN transcriptional factors in naïve vs resistant tumors. (F) The expression of melanocytic markers and Wnt ligands in a human neuroblastoma cohort (TARGET study)^21^. (G) *RBM39* expression from normalized RNA-seq in naïve vs resistant SJNB14 tumors. (H) Volcano plot shows the differential peaks of H3K27Ac in naïve vs resistant tumors by CUT&Tag analysis. (I) IGV plots shows the peak changes of at the loci of *TWIST1*, *PHOX2B* and *S100B*. (J) GSEA shows the upregulation of H3K27Ac at the genomic loci of MES TFs. (K) GSEA shows the downregulation of H3K27Ac at the genomic loci of ADRN signature genes.

Similarly, mice implanted with SIMA (with MYCN amplification) cell line–based xenografts also showed excellent outcome after the completion of a 10-day treatment, but the neuroblastoma eventually relapsed (**Figure S3A**). To test if the relapsed SIMA tumors remained sensitive to indisulam, we resumed treatment after tumor recurrence. Like SJNB14 PDX, SIMA xenografts had 100% complete response after two additional repeated treatment cycles (**Figure S3A**). Western blot analysis of xenografts harvested at 24-hours after the final dosing from the third cycle of treatment confirmed that RBM39 expression was nearly completely abrogated in the treatment group (**Figure S3B**). RNA splicing analysis of tumor xenografts showed that indisulam induced very similar events to those induced by RBM39 knockdown and *in vitro* indisulam treatment, including genome-wide splicing anomalies in skipped exons^30^ (**Figure S3C**). RNA-seq and GSEA analysis revealed that tumor cells displayed upregulated gene signatures of neural crest migration and type I interferons (**Figure S3D**), although the changes in ADRN and MES signatures were not significant. Nevertheless, manual examination of master transcriptional factors of ADRN and MES revealed that the tumors treated with indisulam expressed higher levels of *C-MYC*, *PRRX1*, *FOSL2* but reduced levels of *PHOX2A*, *PHOX2B* and *MYCN* (**Figure S3E**), indicating that cell state underwent a transition from ADRN to MES during the indisulam therapy. Like SJNB14, a significant induction of melanocytic markers and Wnt ligands was observed in the SIMA tumors treated with indisulam (**Figures S3D, 3E**).

Changes in neuroblastoma cell states are associated with epigenetic reprogramming^45,46,48^. To test if this occurred in the indisulam-resistant tumors, we performed Cleavage Under Targets and Tagmentation (CUT&Tag) to map the genome-wide H3K27Ac in SJNB14 naïve vs resistant tumors. H3K27Ac is an epigenetic mark indicating active promoters and enhancers in gene transcription. We identified 8580 differential H3K27Ac peaks (**Figure 2H**). The enrichment of H3K27Ac at *PHOX2B* (a master transcription factor in ADRN) locus was greatly reduced in the resistant tumors, while the H3K27Ac levels were upregulated at the locus of *S100B* (a marker of SCP and MES) (**Figure 2I**). S100B is also a melanoma marker^64^. We also found that H3K27Ac peaks at the *TWIST1* locus were among the top downregulated in resistant tumors (**Figures 2H, 2I**), in line with its reduction in mRNA levels (**Figure 2E**). In neuroblastoma, TWIST1 co-occupies enhancers with MYCN and is required for MYCN– dependent proliferation^65^. SJNB14 is a *MYCN*-amplified tumor. Motif analysis for the H3K27Ac peaks revealed that transcription factors including the MES (JUN-AP1,FOSL2) could bind at the H3K27Ac loci in both naïve and resistant tumors (**Figure S4A**). Then, we performed GSEA analysis for the genes with altered H3K27Ac peaks. Consistent with the RNA-seq results, the indisulam-resistant tumor cells showed a significant enrichment of H3K27Ac at gene loci of MES transcriptional factors (i.e., SMAD3, TEAD4, MYC, **Figures 2J, S4D**), mesenchymal genes such as those involved in muscle contraction, HIPPO (YAP/TAZ) signaling pathway, and interferon alpha and beta signaling (**Figure S4B**). However, there was a significant downregulation of H3K27Ac peaks at the genomic loci of ADRN genes, and those involved in G2/M phase of cell cycle (**Figures 2K, S4C**). These data support that the epigenetic and transcriptional landscapes in the naïve tumors have been reprogrammed when tumor cells developed therapy resistance.

Taken together, these data provided additional evidence that the tumor cells not only switched their state from ADRN to MES, but also acquired additional molecular features such as those expressed in melanomas once they developed resistance, further emphasizing the cellular plasticity of neuroblastoma cells which is reminiscent of cellular pliancy of neural crest or SCP cells.

### Indisulam resistance is associated with RBM39 upregulation and lineage switch from MES to ADRN in a C-MYC–overexpressing xenograft model

Both *Th*-*MYCN/ALK^F1178L^* and the *MYCN-*amplified SJNB14 PDX models demonstrated the cell state switch from ADRN to MES when they developed therapy resistance although a more complex and multi-directional transdifferentiation underlies the de facto mechanism (**Figures 1, 2**). To some degree, our data are in line with the hypothesis that the conversion of ADRN to MES may account for the chemoresistance of neuroblastoma cells. However, the conversion of MES to ADRN of neuroblastoma cells in therapy resistance is less clear. To explore whether the neuroblastoma cells in an MES state could convert to an ADRN state under therapy selection, we treated SK-N-AS xenografts (C-MYC overexpression by a translocated super-enhancer^13^, known as MES dominant^45,46^) with indisulam. Again, SK-N-AS xenografts also showed 100% tumor regression after a 10-day dosing with 25 mg/kg of indisulam (**Figure S5A**) and remained sensitive to indisulam until the 4^th^ repeated treatment cycle. RNA-seq and GSEA analysis revealed that indeed the resistant tumor cells acquired ADRN and neuronal gene signatures with a significant reduction of MES and interferon signatures (**Figures 3A-E**), supporting the hypothesis of interconversion of ADRN and MES states of neuroblastoma cells. Like the *MYCN/ALK^F1178L^* resistant tumors, *RBM39* mRNA was increased by about 4-fold in the resistant SK-N-AS xenografts (**Figure 3F**), but not its paralog, *RBM23* (**Figure 3F**), which has previously been shown to have no effect on RNA splicing^31^. These data further verified that RBM39 was the specific target of indisulam that was responsible for the therapeutic efficacy, leading to a selective pressure on RBM39 in resistant tumors. At the same time, cells underwent cell state alterations from MES to ADRN.

**Figure 3.**
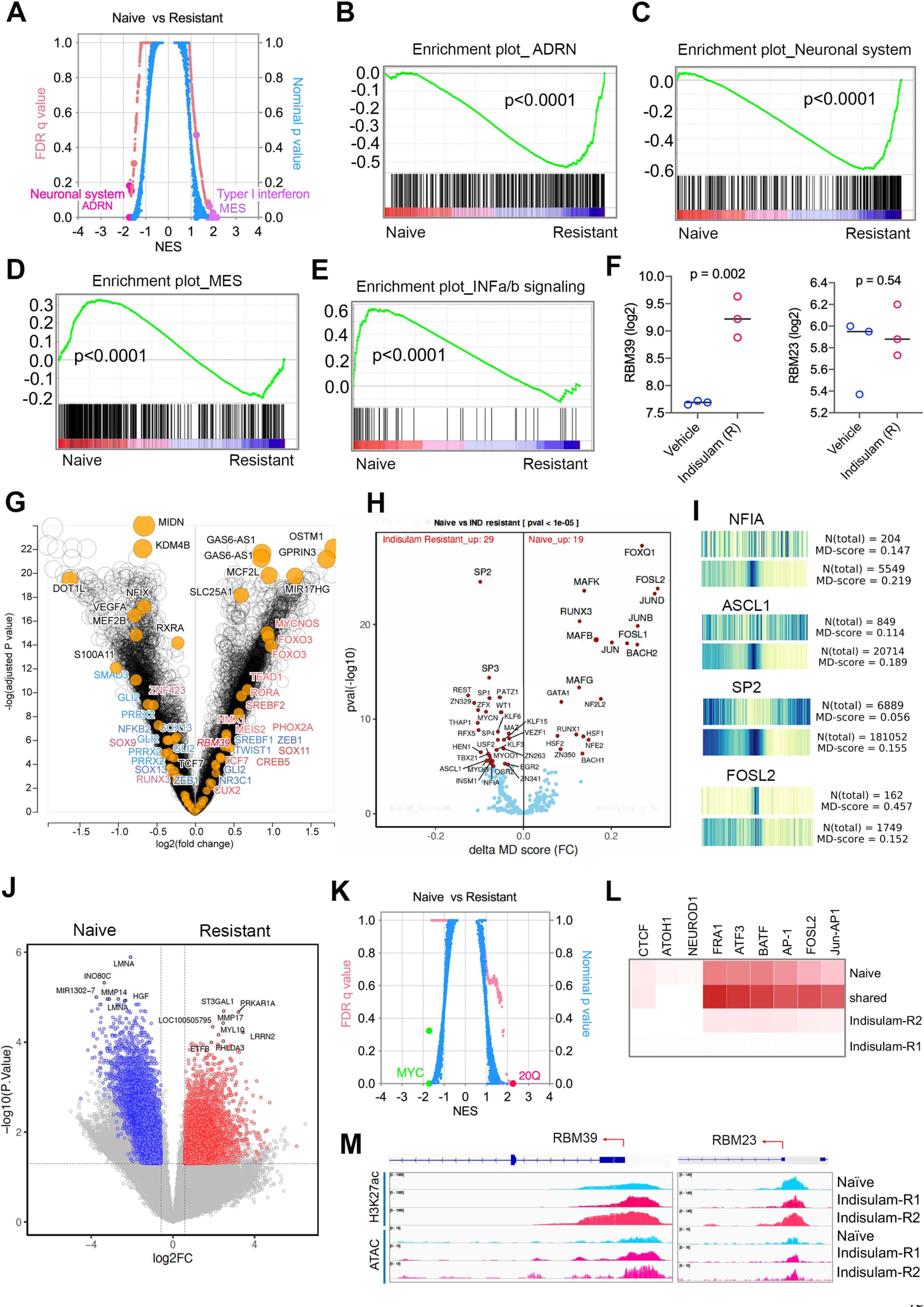
Indisulam resistance is associated with RBM39 upregulation and lineage switch from MES to ADRN in a *C-MYC*-overactive xenograft model. (A) Plot shows GSEA analysis results of RNA-seq data of naïve tumors vs relapsed tumors. Blue color indicates nominal P values and pink color indicates false discovery rate (FDR) q values. (B-E) GSEA plots shows “ADRN” and “neuronal system” gene sets are significantly upregulated in resistant tumors while the “MES” and “Interferon” gene sets are significantly downregulated. (F) *RBM39* and *RBM23* expression from normalized RNA-seq in naïve vs resistant SK-N-AS tumors. (G) Volcano plot shows the differential peaks of ATAC-seq on or at nearby genes in naïve vs resistant tumors. (H) Volcano plot shows the differential binding motifs of transcriptional factors predicted from peaks of ATAC-seq in naïve vs resistant tumors. (I) Heatmap showing the motif displacement (MD) distribution of NFIA, ASCL1, SP2 and FOSL2 (increasingly dark blue indicates increasing motif frequency), MD-score, and the number of this motif within 1.5 kb of an ATAC-seq peak naïve vs resistant tumors. (J) Volcano plot shows the differential peaks of H3K27Ac in naïve vs resistant tumors by CUT&Tag analysis. (K) Plot shows GSEA analysis for genes with differential peaks of H3K27Ac in naïve vs resistant tumors by CUT&Tag analysis. Blue color indicates nominal P values and pink color indicates false discovery rate (FDR) q values. (L) Heatmap indicates the binding motifs for transcriptional factors enriched in naïve, resistant and both based on Homer Motif analysis of H3K27Ac peaks. (M) Snapshots of using the IGV program displaying the peaks in naïve vs resistant SK-N-AS cells by H3K27Ac CUT&Tag and ATAC-seq at *RBM39* and *RBM23* genomic loci.

### Epigenetic reprogramming of indisulam–resistant c-MYC–overexpressing neuroblastoma cells

We hypothesized that the resistant SK-N-AS neuroblastoma cells had undergone an epigenetic reprogramming to alter their MES cell state and thus enhancing the transcription of *RBM39*. To test this hypothesis, we derived cell lines from three independent SK-N-AS indisulam-resistant tumors. These cell lines indeed expressed high levels of RBM39 and were resistant to indisulam treatment in vitro (**Figures S5B,C**). To understand the mechanism of cell state switching, we performed Assay for Transposase-Accessible Chromatin with high-throughput sequencing (ATAC-seq) for mapping genome-wide chromatin accessibility of resistant vs naïve cells. Many ADRN genes such as *PHOX2A* and *MYCN* as well as the *RBM39* locus showed increased chromatin accessibility in the resistant cells (**Figure 3G**), in line with the GSEA results from RNA-seq analysis. Motif analysis for the predicted transcriptional factor binding at the genes with altered chromatin accessibility demonstrated that the MES transcriptional factors such as FOSL1, FOSL2, JUNB, JUND were enriched in naïve SK-N-AS cells, but not in the resistant cells (**Figure 3H, I**). However, the ADRN transcriptional factors such as ASCL1 and NF1A were enriched in the resistant cells. Interestingly, we found that the activity of SP transcriptional factor family members SP2 and SP3 were also enriched in the resistant SK-N-AS cells, together with MYCN (**Figures 3G, I**). While it is unclear what the role of SP transcription factors is in neuroblastoma identity, one early study indicates that they are involved in driving expression of *MYCN* in neuroblastoma cells^66^. To further corroborate our ATAC-seq study, we carried out CUT&Tag to assess the global alterations of H3K27Ac in naïve and resistant cells (**Figure 3J**), an epigenetic mark indicating active promoters and enhancers in gene transcription. GSEA analysis of the differential peaks near the annotated genes revealed that the 20q locus (where RBM39 resides) ranked on the top in the resistant cells while the c-MYC targets ranked on top in the naive SK-N-AS cells (**Figure 3K**). Again, motif analysis of the altered H3K27Ac peaks showed that the binding of MES transcriptional factors such as in the FOS and JUN families was specifically reduced in the resistant cells (**Figure 3L**), similar to the results from ATAC-seq. The *RBM39* locus but not the *RBM23* locus showed a greater increase in H3K27Ac peak in the resistant cells, in line with the elevated ATAC-seq peak (**Figure 3M**), which explains why the resistant cells expressed higher levels of RBM39. These data indicate that the resistant SK-N-AS cells acquired a new cellular state under the selection of repeated indisulam treatment through an epigenetic reprogramming.

### Cell state alterations are coupled with switch of dependency on epigenetics and lineage– specific transcription factors

We surmised that the distinct epigenetic landscapes of naïve and resistant tumor cells may result in dependency on specific epigenetic modifiers and lineage–specific transcription factors in these cells, as cell differentiation and de-differentiation is regulated by a cascade of lineage–specific transcriptional factors (TFs) in cooperation with epigenetic modifiers. To test this hypothesis, we performed CRISPR-Cas9 screening (human epigenetic library with 8 gRNAs/gene and transcription factor library with 4 gRNAs/gene) to knock out these genes for a dropout screen with MAGeCK analysis in naïve versus resistant cells (**Figure 4A, Supplemental spreadsheets 1, 2, 4, 5**), in which reduction of gRNA reads (dropout) indicates that cells are dependent on it for survival. In the transcription factor library screening, we identified more pre-mRNA splicing factors in the resistant cells, particularly when they were cultured under indisulam selection (**Figure 4B**), which is consistent with the function of RBM39 being critical to splicing in neuroblastoma^30^, which also verified the screening robustness. Then we specifically examined the ADRN and MES transcriptional factors (**Figure 4C**). While a general transcriptional regulator CDK9 was essential to both naïve and resistant cells (**Figure 4D**), c-MYC and FOSL2 of MES TFs in the naïve cells were ranked above the ADRN TF HAND2, TBX2 and PHOX2B (**Figure 4C, left**). However, in the resistant cells with or without indisulam selection, c-MYC and FOSL2 dropped down while HAND2, PHOX2B and TBX2 became more essential to the survival of the resistant cells (**Figures 4C, middle and right, 4E, 4F**). These data demonstrate that naïve cells and indisulam-resistant cells have specific dependencies on transcriptional factors that determine the cell state of neuroblastoma.

**Figure 4.**
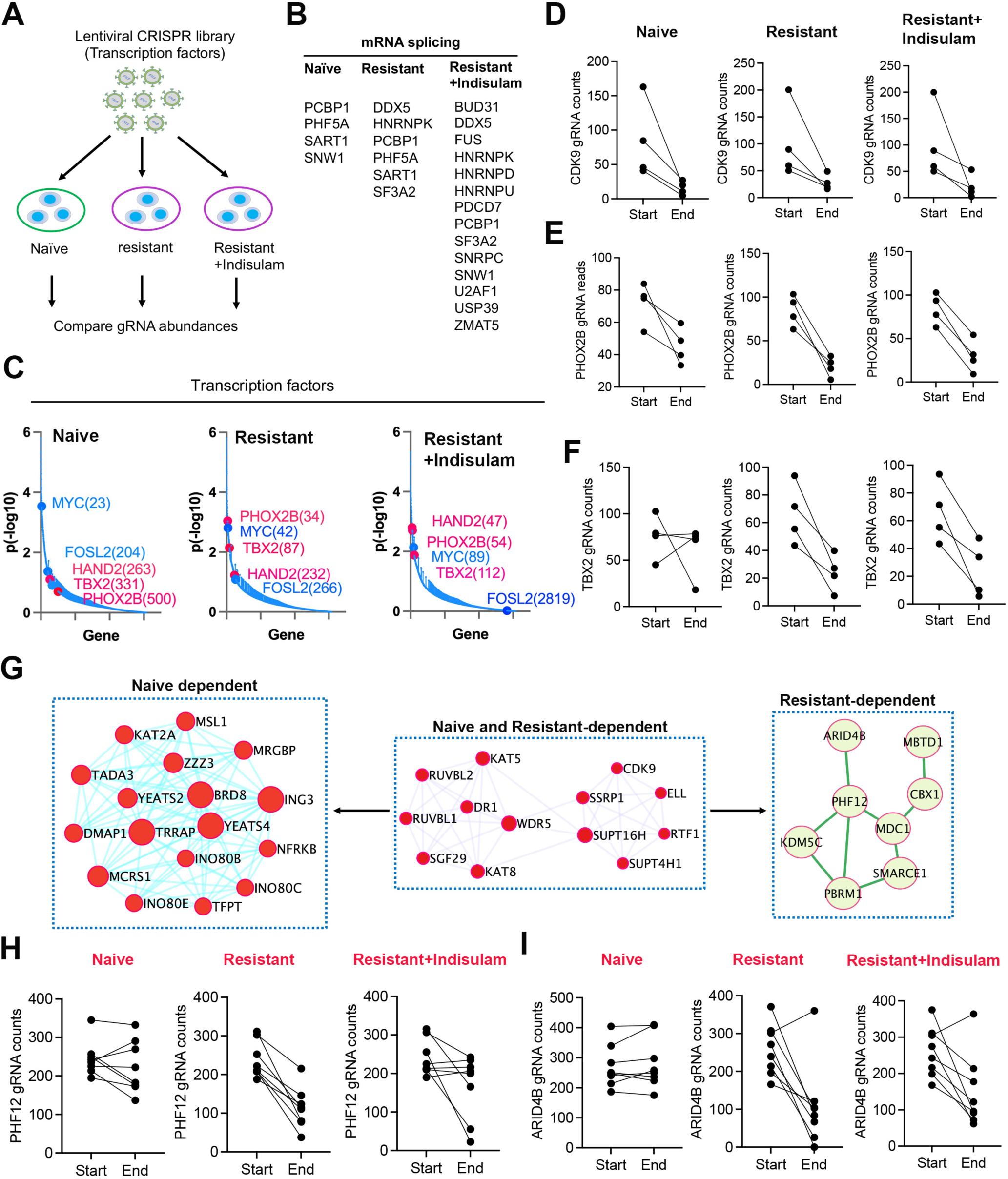
Cell state alterations are coupled with switch of dependency on epigenetics and lineage–specific transcription factors. (A) Focused CRISPR library screening procedure using SK-N-AS cell lines derived from naïve and indisulam-resistant SK-N-AS tumor cells cultured for 3 weeks. (B) Dependency genes related to pre-mRNA splicing in naïve and resistant SK-N-AS cells with or without 250nM of indisulam culture. (C) Ranked dependency genes encode transcriptional factors in naïve and resistant SK-N-AS cells with or without 250nM of indisulam culture. Blue indicates MES TFs, and pink indicates ADRN TFs. (D-F) Normalized counts of gRNAs for CDK9, PHOX2B and TBX2 in naïve and resistant SK-N-AS cells with or without 250nM of indisulam culture. (G) Protein interaction network analysis (STRING: https://string-db.org) of epigenetic regulators that are selectively essential to naïve or indisulam-resistant tumors. (H, I) Normalized counts of gRNAs for PHF12 and ARID4B in naïve and resistant SK-N-AS cells with or without 250nM of indisulam culture.

Then, we compared the epigenetic dependency of naïve and resistant cells, in which the epigenetic modifiers formed unique protein-protein interaction network modules (**Figure 4G**), consistent with the fact that epigenetic modifiers usually form protein complexes to function. We identified chromatin remodeling INO80 complex was more critical to the survival of naïve SK-N-AS cells while the SWI/SNF complex components (PBRM1, SMARCE1) and SIN3A complex components (PHF12, ARID4B) were more important in the resistant cells (**Figures 4G-I**). Notably, we observed that INO80C was one of the top genes with active H3K27Ac in naïve SK-N-AS cells (**Figure 3J**), providing additional evidence that the INO80 complex might be specifically important to the naïve SK-N-AS cells dominated by an MES state while SWI/SNF complex could be more essential to resistant SK-N-AS cells that acquired an ADRN state.

### Selective and shared dependency of naïve and resistant neuroblastoma cells on targetable kinases

The dependency switch of neuroblastoma cells on epigenetic modifiers and lineage–specific TFs suggests that additional dependencies may correspondingly change once neuroblastoma cells convert to another state. As most epigenetic regulators such as ARID4B and transcription factors are difficult to target, we tested if we could identify targetable vulnerabilities of indisulam-resistant neuroblastoma tumor cells by focusing on kinases. Again, with a similar approach, we screened naïve SK-N-AS and indisulam–resistant SK-N-AS cells cultured with or without 250nM indisulam, using the human kinase CRISPR library (**Figure 5A**, **Supplemental spreadsheet 3,** 6 gRNAs/gene). We identified 43, 31 and 25 kinases that were essential for either naïve, or indisulam resistant cells under selective pressure or no selective pressure, respectively; and 13 of them were commonly shared (**Figure 5B, Supplemental spreadsheet 6**). We found that *CLK3*, *DYRK1A* and *DYRK1B*, members of the dual-specificity serine/threonine and tyrosine kinase that play an important role in pre-mRNA splicing by phosphorylating serine- and arginine-rich splicing factors in spliceosomal complex, are essential to naïve SK-N-AS cells but not for the indisulam-resistant cells (**Supplemental spreadsheet 6**). This is in line with the mechanism of indisulam that targets splicing. However, the changes in expression levels of these kinases in resistant vs naïve cells were not correlated with the dependency switch (**Supplemental spreadsheet 7**), suggesting that kinases are not the determinant of cell state. We then specifically examined the targetable kinases with inhibitors available in clinical trials and found 9 kinases are essential to SK-N-AS cells regardless of indisulam resistance, and 2 kinases (*FGFR4* and *CDK2*) are specifically essential to indisulam-resistant SK-N-AS cells (**Figure 5C**). These kinases play critical roles in the G2/M phase cell cycle (i.e., *AURKA*, *AURKB*, *PLK1*), DNA repair (*ATR*, *CHEK1*) and gene transcription (*CDK7*, *CDK9*). We hypothesize that the combination of indisulam with any of these kinase inhibitors may enhance the efficacy and blunt disease relapse. To test this, we chose gartisertib, a selective ATR inhibitor currently in clinical trials^67^, in combination with indisulam with 10 mg/kg, for a two-week treatment. Indeed, gartisertib in combination with indisulam significantly extended mouse survival (**Figure 5D**) and showed very limited toxicity based on the results of body weight gain over time (**Figure 5E**), suggesting that this combination is safe and effective. Recent studies have shown that ATR inhibition is able to reverse the chemoresistance of ALT (alternative lengthening of telomere) neuroblastoma due to telomere dysfunction–induced ATM activation^68^, and significantly enhances the efficacy of ALK inhibition in transgenic neuroblastoma models driven by *MYCN* and *ALK*^69^. These promising pre-clinical data together with our preliminary results provide a rationale to continue testing the combination of indisulam with ATR inhibitors in more mouse models in the future.

**Figure 5.**
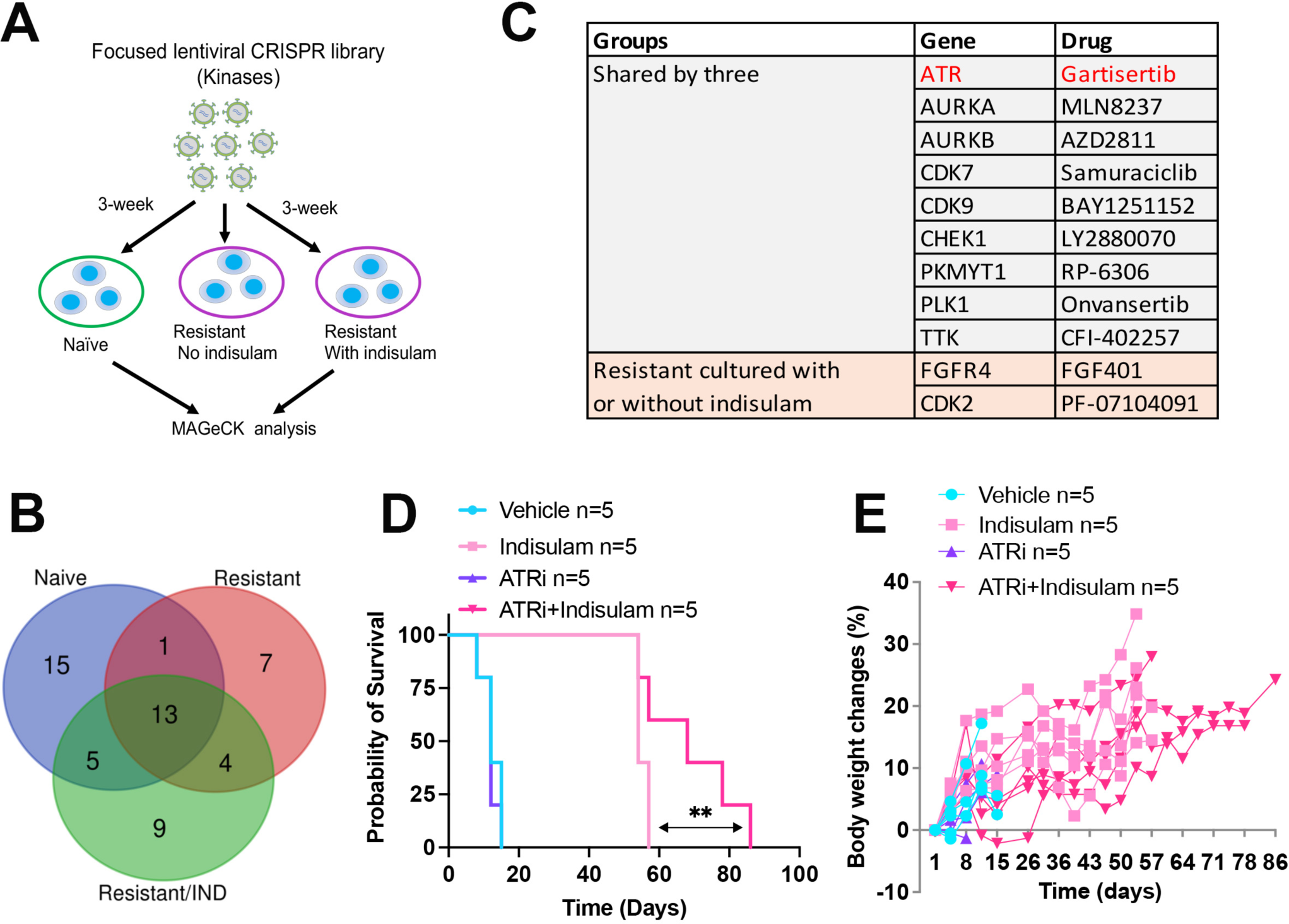
CRISPR identification of targetable essential kinases leads to a more effective combination therapy consisting of indisulam and an ATR inhibitor. (A) Human Kinase CRISPR library screening using SK-N-AS cell lines derived from naïve tumors and indisulam-resistant tumors. (B) Venn diagram showing the shared and cell-type-specific essential kinases. (C) Identified kinases with inhibitors in clinical trials. (D) Kaplan-Meier survival for SK-N-AS tumors treated with vehicle, indisulam, gartisertib, or combination. Indisulam is administered with 10 mg/kg, 5 days on, 2 days off, for 3 weeks, with additional two weeks when tumors relapsed with indisulam alone treatment. Gartisertib is administered with 10 mg/kg, once weekly. ** p = 0.02. (E) Body weight monitoring over time for each individual mouse.

### Lineage switch induced by indisulam treatment can lead to upregulation of GD2 expression through epigenetic reprogramming

GD2 is a disialoganglioside that is biosynthesized from the precursor gangliosides GD3/GM3 by β-1,4-N-acetylgalactosaminyltransferase (B4GALNT1, GD2 synthase) (**Figure 6A**). The expression of GD2 in normal tissues is restricted to the brain, peripheral pain fibers, and skin melanocytes, but it is abundantly expressed in neuroectodermal tumors including neuroblastoma^70^. The application of anti-GD2 immunotherapy has greatly improved the survival of high-risk neuroblastoma patients when it is combined with a differentiating agent and chemotherapy^71–73^. Recent advances in the development of GD2–based CAR T and CAR NKT therapies have provided additional evidence showing the potential of anti-GD2 therapy for patients with relapsed disease^74,75^. While many reasons could account for the response failure of anti-GD2 immunotherapy for a fraction of patients, one of the hypotheses is that lineage switching may lead to low abundance of GD2 expression due to downregulation of GD2 synthase genes. A recent study has shown that cell lines derived from *TH-MYCN* transgenic tumors lost GD2 expression, accompanied with switching of cell state from ADRN to MES^76^. Another study further supported that neuroblastoma state transition to MES confers resistance to anti-GD2 antibody via reduced expression of ST8SIA1^77^. To understand the impact of cell state alterations induced by repeated indisulam treatments on GD2 expression, we examined the key enzymes of GD2 synthesis (*B4GALNT1* and *ST8SIA1*) in our indisulam–resistant models. Our RNA-seq results showed that, in the transgenic *TH-MYCN/ALK^F1178L^*model, one out of two resistant tumors showed a higher expression of *B4GALNT1* and *ST8SIA1* (**Figure 6B**), while the SJNB14 PDX model showed no remarkable changes (**Figure 6C**). In the SIMA model, *ST8SIA1* was significantly upregulated in the tumors with three rounds of indisulam treatment while the expression of *B4GALNT1* showed no changes (**Figure 6D**). In the SK-N-AS model, the expression of both *B4GALNT1* and *ST8SIA1* was significantly upregulated in the resistant tumors (**Figure 6E**), which is in line with the elevated enrichment of H3K27Ac and enhanced chromatin accessibility at the promoter regions of *B4GALNT1* and *ST8SIA1* (**Figures 6F, G**), indicating that epigenetic reprograming of SK-N-AS cells lead to upregulation of GD2 synthesis genes. Then, we applied flow cytometry analysis to profile the GD2 expression of the three resistant cell lines derived from SK-N-AS tumors. The result showed that, in comparison with the naïve control, all three resistant clones expressed higher levels of GD2 (**Figure 6H**). These data indicate that cell lineage alterations induced by indisulam do not lead to reduction of GD2 expression; rather, it results in enhanced expression, at least in some neuroblastomas. These data provide a rationale for a combination therapy of indisulam in combination with anti-GD2 immunotherapy.

**Figure 6.**
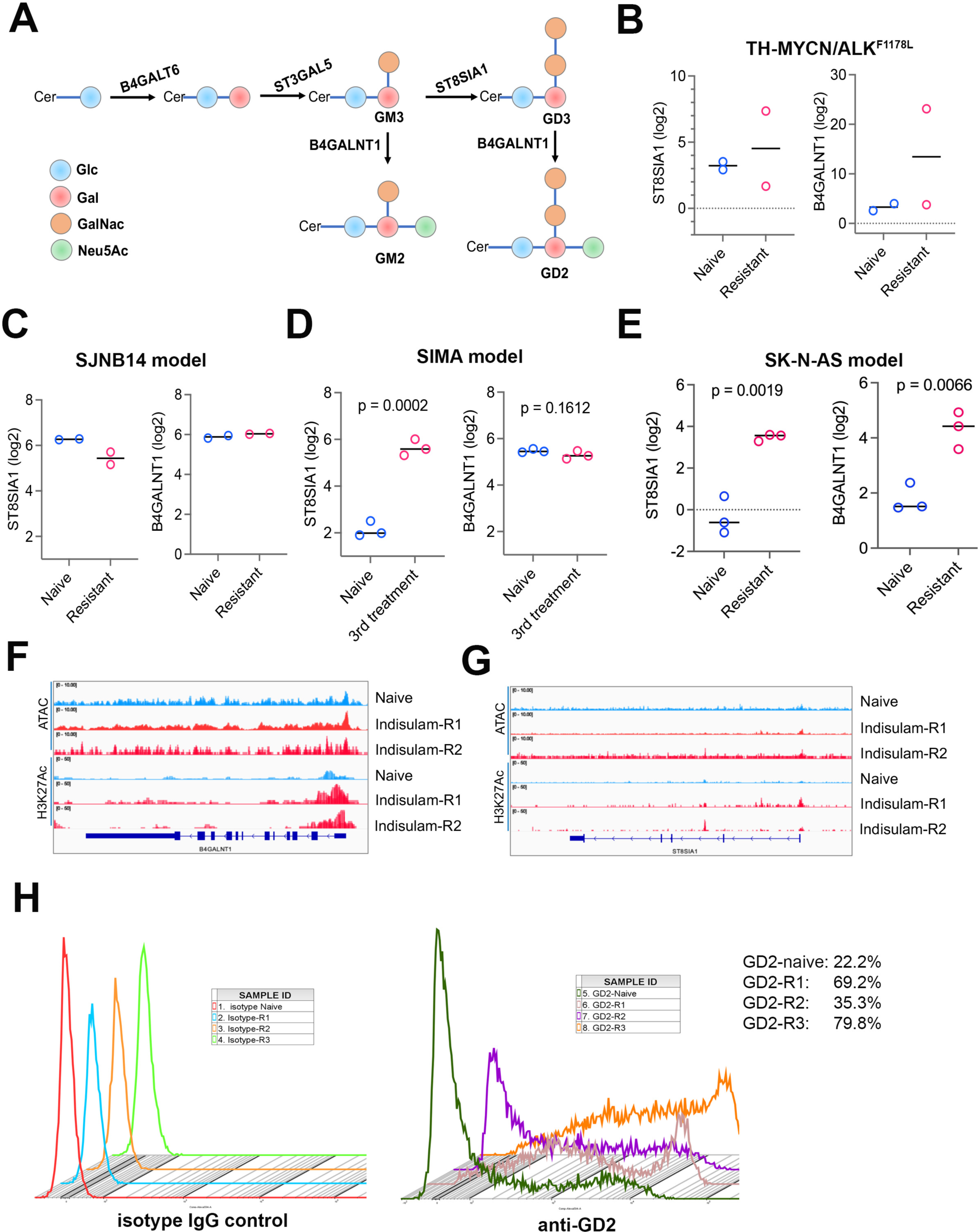
Lineage switch induced by indisulam treatment can lead to upregulation of GD2 expression through epigenetic reprogramming. (A) Cartoon depicts the GD2 synthesis and related key enzymes. Cer, ceramide, N-acylsphingosine; Glc, glucose; Gal, galactose; GalNac, N-acetylgalactosamine; Neu5Ac, N-acetyl neuraminic acid. (B-E) Expression of the two key enzymes in GD2 synthesis in 4 different neuroblastoma models. (F, G) Snapshots of H3K27Ac CUT&Tag and ATAC-seq at *B4GALNT1* and *ST8SIA1* genomic loci using the IGV program displaying the peaks in naïve vs resistant SK-N-AS cells. (H) Flow cytometry analysis of GD2 expression in naïve and indisulam-resistant SK-N-AS cells. Left, isotype IgG as control; right, anti-GD2 intensities.

### Complete eradication of C-MYC driven neuroblastoma in NK cell competent mice

We previously showed that indisulam induced durable complete responses in *C-MYC*– and *MYCN/ALK^F1178L^*–driven neuroblastoma models under immune competent settings^30^. One recent study indicates that indisulam induces neoantigens and elicits anti-tumor immunity, augmenting checkpoint immunotherapy in a manner dependent on host T cells and peptides presented on tumor MHC class I^78^. The excellent efficacy of indisulam in immune competent neuroblastoma models leads to one hypothesis that indisulam modulates T cell immune responses that eliminate cancer cells by making neoantigens through altered pre-mRNA splicing. Thus, despite the cell plasticity of neuroblastoma that may lead to therapy resistance, harnessing the immune system activated by indisulam may lead to a disease cure. To test this hypothesis, we implanted syngeneic *C-MYC*–driven neuroblastoma (derived from *Dbh-iCre/CAG-C-MYC* mouse^30^) into immune competent C57BL/6 mice, Rag2^-/-^ mice (no T and B cells, intact NK cells), and immune deficient NSG mice without T, B and NK cells (**Figure 7A**), which were then treated with indisulam as indicated schedules. Given the high potency of indisulam against neuroblastomas^30^, we began treatment after tumor sizes reached over 1000 mm^3^. Again, we observed exceptional efficacy of indisulam in C57BL/6 mice, which showed complete responses (**Figure 7B**). Interestingly, all tumors in Rag2^-/-^ mice also underwent complete responses (**Figure 7B**, note that the dosing schedule for the two models is one dose per week in Fig. 7B). However, indisulam showed little efficacy to this *C-MYC*–driven mouse neuroblastoma model in immune deficient NSG mice, and progressive disease occurred even given 5 days treatment per week for two weeks. In line with the tumor responses, Kaplan-Meier analysis shows that indisulam treatment led to durable complete responses in both C57BL/6 and Rag2^-/-^ mice (**Figure 7C**). While Rag2^-/-^ mice were sacrificed earlier than the C57BL/6 mice, it was not because of disease relapse rather it was due to mouse aging related illness of this specific strain. These data indicate that it is likely NK cells but not T and B cells that play a critical role in indisulam-mediated anti-cancer activity in neuroblastoma, suggesting that indisulam activates the innate immunity to exert anti-tumor effect. NK cells are known to mount rapid responses to damaged, infected or stressed cells, and play a major role in first-line innate defenses against viral infection and tumor growth^79^. To verify the anti-tumor role of NK cells in Rag2^-/-^ mice, we performed FACS analysis to examine infiltration of the interferon gamma producing NK cells in tumors treated with vehicle and indisulam for 3 days, respectively. Indeed, the NK cell frequency among all cells sorted from the tumors was significantly increased in the indisulam group in comparison with the control group (**Figure 7D**), suggesting that indisulam treatment leads to an inflamed tumor microenvironment with a high percentage of NK cell infiltration. Our data was consistent with a recent report showing that indisulam can induce an inflamed state of acute myeloid leukemia cells^80^, consequently leading to an enhanced NK cell–mediated killing in *in vitro* assays.

**Figure 7.**
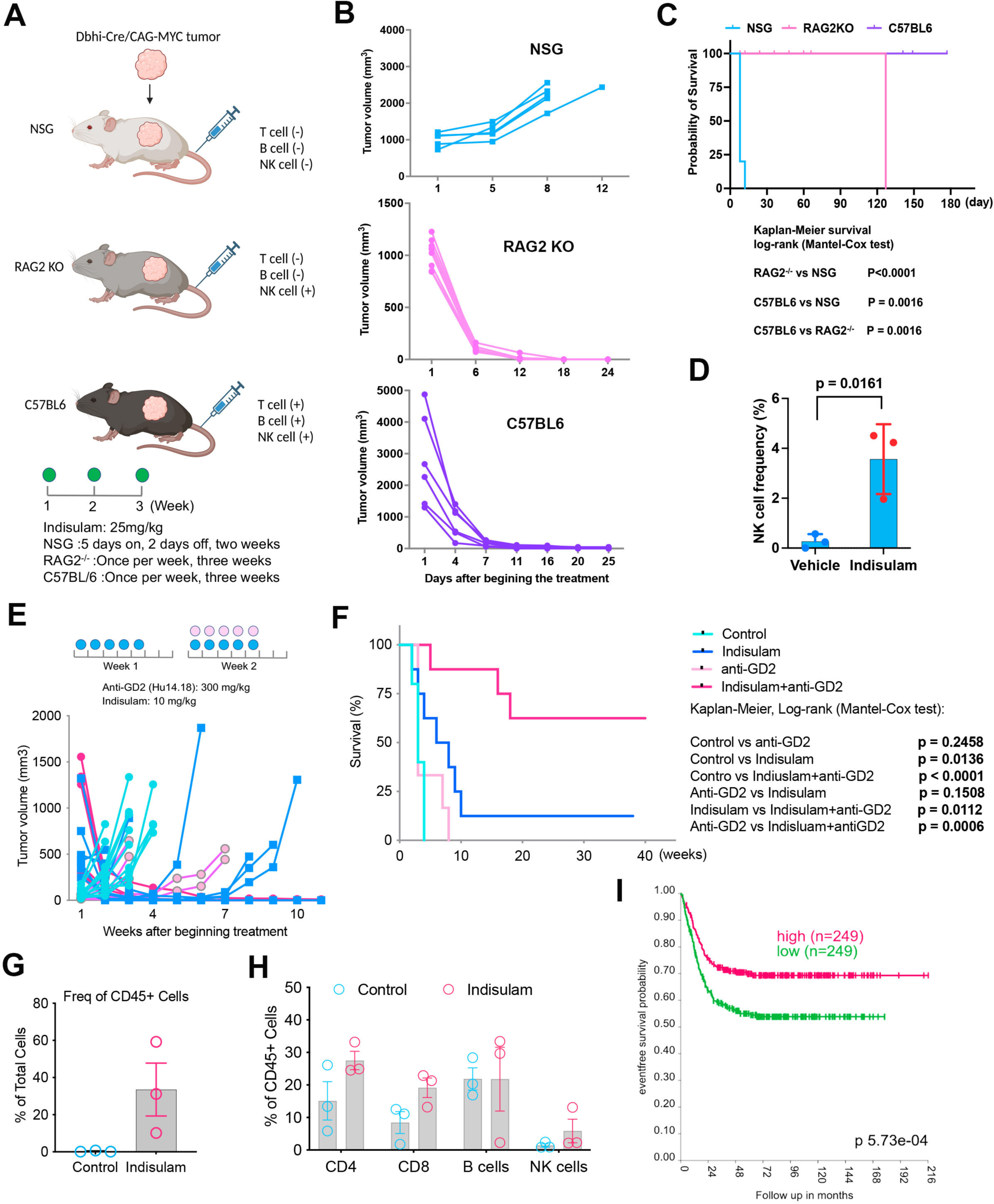
Potent efficacy of indisulam in immune competent mice. (A) Tumors derived from C-MYC transgenic neuroblastoma model implanted into immune deficient NSG mice, T and B cell deficient Rag2^-/^ mice and immune competent C57BL6 mice, with indicated therapeutic schedules. (B) Individual tumor volume for each group treated with indisulam. NSG (n=8), Rag2^-/-^ (n=7), C57BL/6 (n=6). (C) Kaplan-Meier survival curve for each group of mice treated with indisulam, with indicated p values for comparisons of each group. (D) Percentage of NK cells in c-MYC tumors treated with vehicle (n=3) and indisulam (n=3) for 3 days, analyzed by flow cytometry. (E) Individual tumor volume for each group treated with indisulam. TH-MYCN/ALK^F1178L^ mice treated as indicated doses and schedule. (F) Kaplan-Meier survival for transgenic MYCN/ALK^F1178L^ mice treated with vehicle, anti-GD2, indisulam, combination as shown in Figure E, with indicated p values for comparisons of each group. (G) Percentage of immune cells (CD45) in tumors treated with vehicle (n=3) and indisulam (n=3) for 5 days, analyzed by flow cytometry. (H) Percentage of different types of immune cells in tumors treated with vehicle and indisulam for 5 days, analyzed by flow cytometry. (I) Kaplan-Meier survival for neuroblastoma patients with high and low expression of *NCR1* gene in SEQC cohort (GSE62564).

### Durable complete response of anti-GD2 immunotherapy in combination with indisulam against the TH-MYCN/ALK^F1178L^ model

Pediatric solid tumors including neuroblastoma are generally immune “cold”, making it challenging to develop effective immune checkpoint blockade (ICB) therapies, as evidenced by clinical trials^81^. However, the excellent efficacy of indisulam to *C-MYC* tumors in Rag2^-/-^ mice suggests that the innate immunity, including NK cells can be leveraged to develop more effective therapies against neuroblastoma. The application of anti-GD2 immunotherapy has greatly improved the survival of neuroblastoma patients when it is combined with differentiating agents and chemotherapy^71–73^. The anti-GD2 monoclonal antibody exerts an NK cell–dependent ADCC–mediated antitumor effect^57^. We therefore hypothesize that combining indisulam with anti-GD2 immunotherapy may achieve a long-term remission or cure of neuroblastoma. While our dosing schedule (25 mg/kg, 5 days on, two days off, two weeks) used in most cases had achieved remarkable responses in multiple high-risk neuroblastoma models^30^, it was based on the maximally tolerated dose in mice (40mg/kg) in a previous animal study that lacked pharmacokinetic (PK) rationale^82^. We identified and investigated a more clinically-relevant indisulam dose and schedule. Briefly, we evaluated the plasma PK profile of indisulam in normal female CB17/SCID mice, approximately 12 weeks in age. Plasma indisulam was quantified with a qualified liquid chromatography – tandem mass spectrometry (LC-MS/MS) assay. A clinically relevant dose for mice was estimated from unbound plasma PK (**Figure S6**) and exposure – namely, the predicted indisulam average concentration over 5 days (Cavg,120hr). Assuming linear, dose proportional PK, and similar plasma protein binding between species, a clinically relevant dose for mice would range from 6.25 to 12.5 mg/kg Dx5 Q3wks (once daily, 5 days per week for 3 weeks). This regimen would approximate the Cavg,120hr achieved with the clinical 160 mg/m^2^ Dx5 indisulam regimen^83,84^. We then tested the efficacy of indisulam (10 mg/kg) in combination with anti-GD2 mAb (Hu14.18K322A) produced by St Jude GMP, in immune competent *MYCN/ALK^F1178L^*transgenic mice. One of the advantages of the version of anti-GD2 antibody (Hu14.18K322A) is it has been modified to abrogate complement binding, thereby reducing pain caused by the antibody but retaining its clinical efficacy^57^. We dosed mice for week 1 with indisulam (10 mg/kg, Dx5, IP route due to the challenge of IV route for younger mice) surmising it may prime an immune response, followed by combination therapy in week 2 (10 mg/kg of indisulam and 300 μg/mouse of Hu14.18K322A given via IP for Dx5) (**Figure 7E**). The tumor response was monitored by ultrasound imaging of *MYCN/ALK^F1178L^* mice that developed neuroblastoma in abdomen. The results showed that while tumors responded to indisulam with 2-week treatment, eventually they relapsed (**Figure 7E**). Anti-GD2 mAb only led to tumor growth delay in 2 out of 6 mice, suggesting this mouse model is recalcitrant to anti-GD2 mAb therapy. Strikingly, the combination of indisulam with anti-GD2 mAb for only two weeks of treatment led to a durable complete response in all tested mice, even with tumor size over 1.5cm^3^. In the combination group, one mouse died for unknown reasons around week 5, and the rest of mice were culled due to other health conditions but none of them died of disease relapse (**Figure 7F**). These data indicate that the combination of indisulam and anti-GD2 mAb is highly effective against this high-risk neuroblastoma model and may have translational feasibility to human patients. To understand the effect of indisulam on immune cells, we harvested the tumors treated with vehicle control and indisulam for profiling immune cell infiltration in the tumors by FACS analysis with cell type-specific antibodies. In comparison with the control tumors, the data showed that tumors treated with indisulam exhibited a greater enrichment of CD45^+^ cells (an indicator of total immune cells) (0.29±0.35% vs 33.5±24.6%) (**Figure 7G**), in line with the enrichment of CD4 T cells (15.1±10.2% vs 27.5±4.9%), CD8 T cells (8.4±5.8% vs 19.1±5.2%), and NK cells (1.5±0.88% vs 5.8±6.3%) but not B cells (21.86±5.9% vs 21.8±17%) (**Figure 7H**). To corroborate the importance of NK cells in anti-neuroblastoma activity, we examined the association of NK cell infiltration in tumors with patient survival by using the NK cell marker gene *NCR1*. *NCR1* encodes NKp46 that is the major NK cell-activating receptor involved in the elimination of target cells and mediates tumor cell lysis. Indeed, a higher expression level of the NK cell marker *NCR1* was correlated with a better event-free survival and overall survival of neuroblastoma patients (**Figurers 7I, S7A**), regardless of risk and *MYCN* status (**Figures S7B-E**). Taken together, it is likely that it is the innate immunity mediated by NK cells in cooperation with indisulam plays the major role in eradicating the tumor cells. These data provide a rationale to translate the indisulam to clinic in combination with anti-GD2 immunotherapy for high-risk neuroblastoma patients.

## DISCUSSION

The cellular heterogeneity of neuroblastoma cells was observed decades ago^42–44^. Recent studies have defined two major populations (ADRN and MES) of neuroblastoma cells according to their transcriptomic and epigenetic features^45–47^. scRNA-seq analysis of neuroblastomas revealed that ADRN exists in nearly all tumors with variable degrees of MES signature^51,52,54,85,86^. Using our transgenic *MYCN/ALK^F1178L^*model, we found that once tumor cells developed indisulam resistance, they predominantly acquired SCP features seen during normal medulla development^51–53^. SCP has high cellular pliancy with potential to differentiate to different cell lineages depending upon the environmental cues. In an ADRN PDX model (*MYCN* amplified) that developed indisulam resistance, cancer cells lost their ADRN features and were enriched with expression features of melanoma such as MITF. It is well known that the melanocyte is a derivative of neural crest cell or SCP^62,87,88^. Conversely, the MES dominant tumors acquired ADRN features once they developed resistance to indisulam, which was not observed in previous studies. These findings illustrate that neuroblastoma cell states can not only interconvert but could also transdifferentiate to additional lineages from the ADRN and MES states. Thus, our data strongly suggest that the capacity of neuroblastoma cells transdifferentiating to different developmental stages of neural crest progeny is an important mechanism by which neuroblastoma acquires therapy resistance (**Figures 8A, B**).

**Figure 8.**
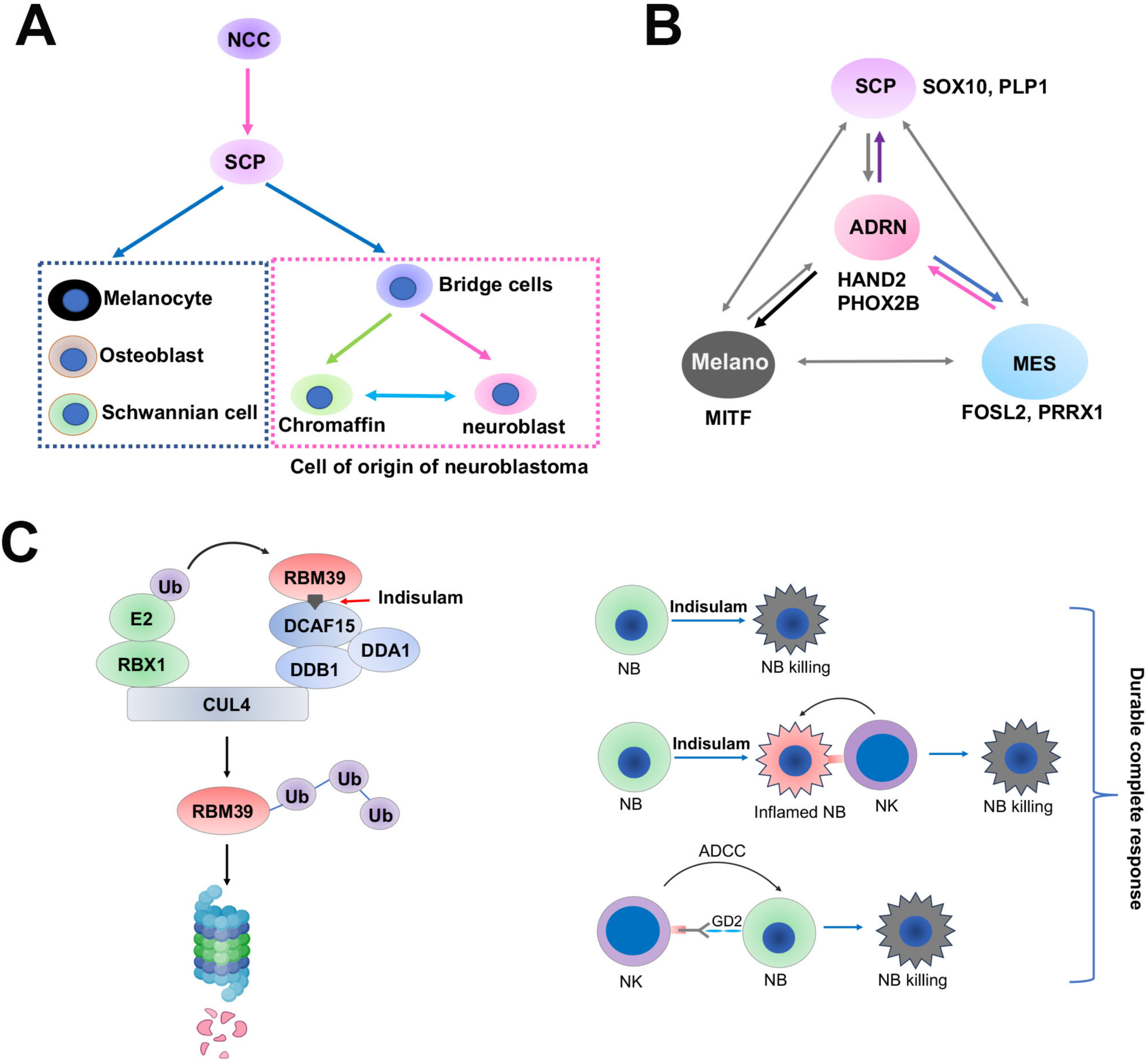
Models for neuroblastoma cell plasticity and working mechanism of indisulam. (A) Schwann cell precursors from neural crest cells could differentiate to distinct lineages including neuroblasts and chromaffin cells that are presumed to be cell of origin of neuroblastoma. NCC, neural crest cell. SCP, Schwann cell precursors. (B) Neuroblastoma cells have multiple cell states that are potentially able to interconvert. These cell states are determined by lineage–specific master TFs or CRC TFs. ADRN, adrenergic. MES, mesenchymal. Melano, melanocytic. (C) left indicates the mechanism of action of indisulam. Right indicates the anticancer mechanism for combination of indisulam with anti-GD2 immunotherapy. Indisulam degrades RBM39 in neuroblastoma cells leading to cell death due to splicing defects (**top**). At the same time, indisulam induces an inflamed state of neuroblastoma cells, leading to NK cell-mediated killing (**middle**). When GD2 immunotherapy is applied, NK cells elicit antibody-dependent cellular cytotoxicity (ADCC) to kill neuroblastoma cells (**bottom**). All three mechanisms together lead to a durable complete response.

Cancers bear transcriptional features of their original tissue lineages under normal developmental programs, and such features are largely determined by a small number of master transcription factors (TFs)^89^. These master TFs typically regulate their own genes through an autoregulatory loop that forms the core transcriptional regulatory circuitry (CRC) of a cell, and their expression is dominated by super-enhancers that drive high rate of transcription^89^. The transcriptional programs of ADRN and MES neuroblastoma cells are dominated by their respective CRC TFs (**Figure 8B**). Our study showed that neuroblastoma cells also switch CRC TF dependency once their cell states change. For example, we noticed that PHOX2B and TBX2, the two ADRN CRC TFs, were less important to the survival of naïve SK-N-AS cells (MES state) but more essential to the indisulam-resistant SK-N-AS cells (ADRN state). The naïve and resistant cells also exhibited distinct dependency on epigenetic modifiers. We found that the naive SK-N-AS cells are more dependent on INO80 chromatin remodeling complex, while the resistant cells were more dependent on the components of SWI/SNF complex (PBAF complex) including PBRM1 and SMARCE1. INO80 and SWI/SNF complexes are ATP-dependent chromatin remodelers. While there is no information about the functions of INO80 complex in neuroblastoma, a previous study has shown that INO80 occupies >90% of super-enhancers in melanoma, and its occupancy is dependent on transcription factors such as MITF^90^. However, a recent study has shown that the expression of SWI/SNF is regulated by SOX11 in neuroblastoma, and SOX11 is an ADRN CRC TF serving as a dependency transcription factor in adrenergic neuroblastoma. In SJNB14 model, the expression of *SOX11* was reduced after the tumors switched their lineage from ADRN to other cell states such as melanoma-like, which conversely expressed high levels of *MITF*. One study shows that SMARCE1 promotes neuroblastoma tumorigenesis through assisting MYCN-mediated transcriptional activation^91^. The components of SIN3A complex (ARID4B, PHF12) also appeared to be important to survival of indisulam–resistant SK-N-AS cells. While the information of ARID4B and PHF12 in cancer, particularly in neuroblastoma, was scarce, PHF12 is a selective dependency gene of neuroblastoma cells (**Figure S8**). Overall, these data mutually support that cell state switching is accompanied by the corresponding alterations of epigenetic and CRC TF dependency. This was also reflected by the distinct dependency of kinases for survival of naïve and indisulam–resistant SK-N-AS models, suggesting that therapies could potentially be developed to block the cell lineage switching to overcome therapy resistance. In the future, how epigenetic modifiers and CRC TFs cooperate to determine the cell state under different cellular context needs to be elucidated.

RNA-seq analysis reveals 27 immune cell subtypes in neuroblastomas, including distinct subpopulations of myeloid, NK, B, and T cells^92^. However, the biological functions of these immune cells in neuroblastoma are not well understood. Recent studies have shown that tumor regression induced by C-MYC loss appears to be NK cell–dependent^93^. *MYCN* is also an immunosuppressive oncogene that dampens the expression of ligands for NK-cell–activating receptors in human high-risk neuroblastoma^94^. Therefore, NK cell–mediated immune surveillance may be critical to constrain neuroblastoma. The application of anti-GD2 antibody in neuroblastoma treatment significantly improves survival of patients with high-risk disease, and anti-GD2 mainly acts through NK cell–mediated antibody–dependent cellular cytotoxicity (ADCC)^71,95,96^. However, not all high-risk patients respond to anti-GD2 therapy. Thus, improving NK–mediated cancer killing for MYC–driven neuroblastoma is attractive. Interestingly, we found that the expression of GD2 in MES type cell was greatly upregulated once they switched to an ADRN state, suggesting that indisulam may potentially enhance the efficacy of anti-GD2 immunotherapy. While indisulam may be important to stimulate the activity of CD8 T cells to boost the efficacy of ICBs by producing neoantigens through alternating pre-mRNA splicing^97^, our study demonstrated that NK cells might be more critical for eliminating cancer cells in our syngeneic *C-MYC* and transgenic *TH-MYCN/ALK^F1178L^* neuroblastoma models. We found that indisulam treatment promoted a remarkable infiltration of NK cells in the c-MYC tumors implanted in Rag2^-/-^ mice, and various immune cells including T and NK cells in the transgenic *TH-MYCN/ALK^F1178L^* tumors, suggesting that indisulam induced an “inflamed” tumor microenvironment. Although it awaits future studies to dissect the functions of each type of immune cells including the CD8 T cells in indisulam–mediated anticancer activity, the durable complete responses induced by combination of anti-GD2 immunotherapy with indisulam suggests that NK cells might play an important role, which was further supported by the C-MYC syngeneic model in that tumors were completely eliminated in NK cell intact Rag2^-/-^ mice that lack functional T and B cells, but little effect was observed in NK/T/B cell deficient NSG mice. One recent study showed that modulation of RNA splicing by indisulam enhances NK cell-mediated tumor surveillance functions^80^. RBM39 in NK cells could possibly affect the anti-tumor functions of NK cells, which needs to be tested in the future studies by using conditional *RBM39* knockout mouse. Overall, our data support that indisulam exerts its anticancer mechanism through three modes (**Figure 8C**). First, indisulam degrades RBM39 in neuroblastoma cells leading to cell death due to splicing defects, which is a major mechanism of indisulam in an immune deficient setting. At the same time, indisulam induces an inflamed state in neuroblastoma tumors, leading to infiltration of immune cells and NK cell–mediated killing in immune competent or innate immunity competent settings. And last, when GD2 immunotherapy is applied, NK cells elicit antibody-dependent cellular cytotoxicity (ADCC) to further enhance the killing of neuroblastoma cells. All three mechanisms together lead to a durable complete response in immune competent neuroblastoma models. Our study also indicates that enhancing NK cell–mediated anti-tumor effect by indisulam could be applied in patients with other cancer types who receive antibody immunotherapy that elicits ADCC.

## Method

### Cell lines and reagents

SIMA (DSMZ, ACC164), SK-N-AS (ATCC, CRL-2137) naive and indisulam-resistant cell lines (R1-R3) were plated in RPMI 1640 (Corning, cat#: 10-040-CM) complete media (10% FBS, Gibco, cat#A5256701, & 1% Pen Strep, cat#:15140-122). All cells were maintained at 37 °C in an atmosphere of 5% CO_2_. All human-derived cell lines were validated by short tandem repeat (STR) profiling using PowerPlex® 16 HS System (Promega) once a month. Once a month a polymerase chain reaction (PCR)-based method was used to screen for mycoplasma employing the LookOut® Mycoplasma PCR Detection Kit (MP0035, Sigma-Aldrich) and JumpStart™ Taq DNA Polymerase (D9307, Sigma-Aldrich) to ensure cells were free of mycoplasma contamination.

Indisulam, obtained from MedKoo Biosciences (catalog #201540), was prepared as a stock concentration of 300mM in DMSO. The process involved rigorous vortexing and sonication until achieving thorough mixing, resulting in a dark red solution. Indisulam was then formulated in 3.5% DMSO, 6.5% Tween 80 (Sigma, P1892-10mL), and 90% saline (Sigma, S8776-100mL), with a sequential addition of chemicals beginning with the indisulam in DMSO, followed by Tween-80, and finally saline. Vehicle solution for a 20g mouse is composed of 7μl DMSO, 13 μl Tween-80 and 180 μl Saline. Indisulam solution for a 20g mouse is composed of 4.3 μl Indisulam (from the 300mM stock), 2.7 μl DMSO, 13 μl Tween-80 and 180 μl Saline, which leads to a final concentration of 0.5 mg/200mL for 25 mg/kg dose. Additionally, the Indisulam solution was further diluted for treatment at 10 mg/kg. In that case, Tween-80 and Saline were employed.

Gartisertib was purchase from MedChemExpress (Cat#HY-136270). To make a 2 mg/ml stock solution of gartisertib, it was suspended in 15% Captisol (Cat#HY-17031)/0.1M hydrochloride acid (LabChem, LC152204).

### Hu 14.18 K322A antibody production

The seed train for the 5L bioreactor was initiated from a single vial of the Master Cell Bank (Hu14.18 K322A YB2/0 CHIL-02 Ab Clone 134 Master Cell Bank) that was thawed and the cells placed into a 125 mL Erlenmeyer flask with cHSFM (HSFM + 2 g/L Soytone + 2 g/L Phytone™ + 4 mM GlutaMax™+ 1000 ng/mL Methotrexate (MTX)) at a cell density of 0.4 x 10^6^ cells/mL and placed into a shaking CO_2_ incubator set at 37°C, 70% humidity and 8% CO_2_ and 120 rpm. Cells are spilt every 48-72 hr reaching six 1000 mL Erlenmeyer flasks (200 mL volume) (Passage 9) providing cells to seed 5L of cHSFM medium at 0.1 x 10^6^ cells/ml in a 50 L WAVE bioreactor. The WAVE bioreactor is cultured until Day 4 post inoculation, with a minimum cell density of 1.00 x 10^6^ cells/mL was achieved. The Hu14.18 K332A cells from the Wave bioreactor were used to seed a starting volume of 2.3 L at 0.25 x 10^6^ cells/mL in the Sartorius Stedim Biotech BIOSTAT 5 L glass bioreactors.

Engineering runs were set up in 5L Sartorius bioreactors with initial charge of 2.4 L of initial charge of cells and HSFM media with 6mM glutamax and 2g/L of Soytone and Phytone. The agitation rate was set at 350 RPM, the pH was set at 6.9, and the dissolved oxygen setpoint was 50%. The cells concentration at time of inoculation was 0.4 x 10^6^ cells /ml. The bioreactors were fed using HSFM containing 6mM glutamax and 10g/L Soytone-Phytone at a constant flow rate starting after 72h post inoculation and continued for 10 days. The cells were harvested when the viability dropped below 50%.

Hu 14.18 K322A culture from a 5 L bioreactor was clarified by centrifugation and filtered using a 0.2 µ nominal filter and a 0.22 µ sterile filter. The filtered supernatant was stored at 4°C until purification. The Hu14.18K322A Ab was purified using MabSelect Sure Protein A resin (Cytiva) equilibrate with PBS. The antibody was eluted from the column using 0.2 M Glycine, pH 3.5. The MabSelect purified Hu14.18 K322A Ab was subjected to a low pH hold step at pH 3.7 for 30 min. The antibody was neutralized to pH 6.0 using 50 mM Malonate buffer and 1M Tris-HCl, pH 8.0. The Hu14.18 K322A Ab was then purified using Capto SP ImpRes resin (Cytiva) equilibrated with 50 mM Malonate buffer, pH 6.0. The Hu14.18 K322A Ab was eluted from the column using 50 mM Malonate buffer, pH 6.0 plus 200 mM NaCL. The Hu14.18 K322A Ab was passed through a Sartobind Q membrane (Sartorius) to remove potential Host DNA and endotoxins. Hu14.18 K322A Ab that was filtered through Sartobind Q membrane filter was diafiltrated into PBS, pH 6.0, 100 mM Arginine and concentrated to approximately 10 mg/ml using a 30 kD Sartocon Slice 200 (Sartorius) membrane. To the concentrated Hu 14.18 K322A Ab, PBS pH 6, 100 mM Arginine,10 % Tween 80 buffer was added so that the final concentration of Tween 80 in the protein was 0.03%. The Hu 14.18 K322A Ab in formulation buffer was sterile filtered through a 0.22 µm Stericup filter (Millipore) and stored at 4°C.

### Western blot

Cells were washed with 1x PBS pH=7.4 while keeping the plate on ice. Removal of 1X PBS is followed by addition of 2x sample loading buffer [0.1M Tris HCl (pH 6.8), 200mM dithiothreitol, 0.01% bromophenol blue, 4% SDS, and 20% glycerol] and scrapped from 6 well plate into a 1.5mL Eppendorf tubes. Keeping the samples on ice, cell lysates were sonicated once with 30% amplitude output (sonics, VIBRA cell) for 5-10 seconds, followed by 10-minute heating at 95°C. Cell lysates are than centrifuged at 1000 rpm for 2 minutes at room temperature before adding 10µL of lysates into 4-15% Mini-Protean TGX stain free gel (BioRad, cat #: 4568086) and transferred into a methanol-soaked polyvinylidene difluoride membrane from (BioRad, cat# L002043B). Membrane is blocked using non-fat dry milk mixed with PBS supplemented with 0.1% Tween 20 (PBST) for one hour while on gentle shaking. Membrane is than washed with PBST for 15 minutes at room temperature, changing the wash solution every 5 minutes. Membrane is than incubated with RBM39 primary antibody (Sigma, cat#:HPA001591, RRID: AB_1079749;1:1000) and HSP90 Primary antibody (Santa Cruz: Cat#:SC13119, (1;1000 dilution, RRID#AB_675659) at 4°C overnight with gentle shaking. The following day, primary antibody is removed and washed for 5-minute intervals of PBST for a total of 15 minutes before adding anti-mouse or rabbit horseradish peroxidase-conjugated secondary antibody (Invitrogen: Ref: A24531 GTXRB IGG F(AB)’2 HRP XADS: 1:4000) mixed with blocking buffer. Membrane is incubated for 1hr at room temperature with gentle shaking and washed with three 5-minute washes with PBST at room temperature. Lastly, the membranes were incubated for 2 minutes at room temperature with a 1:1 of Super Signal West Pico PLUS luminol/ enhancer [ThermoScientific: cat#: 1863096] and stable peroxide [ThermoScientific: cat#1863097] before visualizing antigen-antibody complexes on the Odyssey Fc Imaging System (serial #: OFC. 1358).

### Derivation of indisulam-resistant SK-N-AS cells from indisulam-resistant SK-N-AS tumors

The vehicle-treated SK-N-AS tumor (*n* = 3) and indisulam-resistant SK-N-AS tumors (*n* = 3) were excised and placed in a sterile tube containing phosphate-buffered saline (PBS) on wet ice during transport from the animal research facility to the research laboratory. Tumor samples were manually minced using a sterile scalpel and underwent an enzymatic digestion with collagenase IV (2 mg/ml; in 25 ml of RPMI medium) for 1 hour in a 37°C rotor (Robbins Scientific Corporation, model 2000). After digestion, cells were filtered using a 70-μm sterile strainer and cultured in RPMI medium with 10% FBS and 1% penicillin and streptomycin.

### Cell viability assay

Cell viability for SK-N-AS naive and Indisulam-resistant cell lines (R1-R3) was determined by PrestoBlue cell viability reagent [Invitrogen, Cat#: A13262]. Following the manufacturer’s instructions, cells were plated in two white 96-well plates (PerkinElmer cat#6005680). The first plate contained 90µL of cells and served as a control plate before treatment is added. 10µL of PrestoBlue reagent was added into the cells to assess baseline activity. The second plate followed a 5-day treatment to assess IC_50_. Cells were plated in 45µL of 3000 cells/well and left to incubate at 37°C with 5% CO_2_ for 24 hours. The following day, cells are treated with serially diluted 45µL of 2x concentrations of indisulam until the final concentration within the well resulted in: 0nM, 0.16nM, 0.5nM, 1.5nM, 4.5nM,13.7nM, 41nM,123nM, 370nM,1.1µM, 3.3µM, 10µM. Each concentration is tested in 8 replicates. After 5-day treatment, 10µL of 10X PrestoBlue reagent is added to plates and incubated at 37°C with 5%CO_2_ for 30 minutes. Fluorescence is read at 560-nm excitation/590-nm emission using Synergy H1 microplate reader. Florescent values were normalized to vehicle and graphed on PRISM9 GraphPad software. Normalized data is transformed into log10 and analyzed using nonlinear regression log(inhibitor) vs response (three parameters).

### GD2 flow cytometry analysis

The naïve and indisulam resistant SK-N-AS cells were harvested as a single-cell suspension, washed with PBS, and blocked with blocking buffer. Subsequently, cells were incubated with anti-GD2 (Sigma Aldrich. Catalog # MAB2052) or Isotype control mAb (Thermo Fisher Scientific, Catalog # 14-4724-85) (1 μg per 10^6^ cell) master mix for 1 hour on ice. Following this, cells were washed with PBS supplemented with 5% FBS. Next, the cells were incubated with Alexa Fluor 594-conjugated anti-mouse IgG (1:1000) for 60 minutes in the dark. After three washes, cells were spun down and resuspended in PBS for analysis. The experiment was acquired using BD Fortessa (San Jose, CA). Alexa Fluor 594 signals were detected with a 582/15 band-pass filter under a 562 nm laser excitation. Signal overlay analysis was conducted using FlowJo software for flow cytometry data.

### Sanger DNA sequencing of RBM39

Total RNA was isolated from tissue samples as per the manufacturer’s instructions of the RNeasy Mini Kit (Qiagen, catalog no. 74106). cDNA was prepared from 1 μg of total RNA using a cDNA synthesis kit according to the manufacturer’s protocol (Thermo Fisher Scientific, catalog no. 18091050). RBM39 gene was amplified by PCR using the cDNA as the template along with the following primers (forward: 5′-atttctagagccaccatggcagacgatattgatattgaagcaatgc-3′; reverse: 5′-attggatcctcatcgtctacttggaaccagtagc-3′) using the Phusion High Fidelity PCR Kit (New England Bioloabs, cat#E0553S). The resultant PCR product was gel-purified using a Qiagen gel extraction kit (Qiagen, cat#28706) according to the manufacturer’s protocol. The DNA was sent for Sanger sequencing at Hartwell Center in St. Jude using the following primers: primer 288F, ACAGAAGTCCTTACTCCGGACC; primer 388R, ACTTTTGCTTCGGGAACGTCG; primer 602F, GTCGATGTTAGCTCAGTGCCTC. Primer 602F is used to detect the RBM39 mutation in the RRM2 motif.

### CRISPR library construction, screening and analysis

The customized gRNA libraries (epigenetics, transcription factors, and kinases, Supplemental spreadsheets 1-3) were constructed by following the published protocol^98^. The 20bp gRNAs prepended with extra sequences (TATCTTGTGGAAAGGACGAAACACCG for 5’ and GTTTTAGAGCTAGAAATAGCAAGTTAAAAT for 3’) were synthesized by GenScrip using GenTitan™ Oligo Pool service. Library amplification and Gibson Assembly into the pLentiguide-Puro backbone (Addgene #52963) was performed as previously described^99^. The plasmid library was amplified and validated in the Center for Advanced Genome Engineering at St. Jude Children’s Research Hospital as described in the Broad GPP protocol (https://portals.broadinstitute.org/gpp/public/resources/protocols) except EnduraTM DUOs (Lucigen) electrocompetent cells were used for the transformation step. NGS sequencing was performed in the Hartwell Center Genome Sequencing Facility at St. Jude Children’s Research Hospital. Single-end, 100-cycle sequencing was performed on a NovaSeq 6000 (Illumina). Validation to check gRNA presence and representation was performed using calc_auc_v1.1.py (https://github.com/mhegde/) and count_spacers.py, with qualities passing criteria: Percentage of guides that matched was 82.1%, 80.1% and 80.3%, respectively for epigenetic, kinase and transcriptional factor library (recommended >70%); Percentage of undetected guides was 0 for all three (recommended <0.05); Skew ratio of top 10% to bottom 10% was 1.56, 1.86, 2.66 respectively for epigenetic, kinase and transcriptional factor library (recommended <10); Area under Curve was 0.56, 0.57, 0.6 respectively for epigenetic, kinase and transcriptional factor library (recommended <0.7). For the lentivirus production of CRIPSR libraries, we used PEI-PRO (Polyplus, Cat #115-100) to transfect human embryonic kidney (HEK) 293T cells with CRISPR-gRNA pool library constructs and accompanying helper plasmids: pCAG-kGP1-1R, pCAG4-RTR2, and pHDM-G. The following day, the plates were rinsed, and fresh medium was provided. Over a 72-hour period post-transfection, we gathered replication-deficient lentiviral particles every 12 hours. These particles were concentrated through centrifugation at 28000 rpm, followed by reconstitution in RPMI medium and subsequent storage at -80°C in small aliquots. Estimations for the MOI were made for each library, aiming for an approximate ∼0.3 MOI. To achieve this, cells were transduced with the pooled CRISPR library in the presence of polybrene (8 μg/ml). After 24 hours, cells underwent selection using puromycin (2 μg/ml) for an additional 48-hour duration to enable selection. Subsequently, cell counting was conducted, and live cells were quantified to determine the MOI.

SK-N-AS-Naïve and SK-N-AS-indisulam resistant cell (SKNAS-R2) derived from tumors underwent transduction utilizing Cas9 lentivirus (Addgene, Cat #52962) followed by a selection process employing blasticidin at a concentration of 10 µg/ml (Sigma, Cat #203350) for 7 days. Validation of Cas9 protein expression was performed via western blot analysis. Cas9 expressing SK-N-AS and SKNAS-R2 were transduced with the customized human CRISPR Knockout pooled library, known as epigenetic library, housing 4032 unique sgRNA sequences targeting 480 human genes (including 8 sgRNAs per gene and 192 non-targeting controls), or transcription factor library, housing 10744 unique sgRNA sequences targeting 2558 human genes (including 4 sgRNAs per gene and 512 non-targeting controls) or Kinase library, housing 4032 unique sgRNA sequences targeting 480 human genes (including 6 sgRNAs per gene and 192 non-targeting controls). To ensure effective barcoding of individual cells, we maintained a low MOI (∼0.3). Post-transduction, cells underwent puromycin selection (2 μg/mL, Millipore Sigma) for 48 hours, followed by removal of dead cell debris and maintenance in complete medium. The transduced cells were subjected to treatment with both vehicle and indisulam 250 nM, with concentrations chosen based on PrestoBlue assay (250 nM that show minimal effect on cell proliferation in resistant cells). A minimum of 5×10^6^ cells for the transcription factor library and 3×10^6^ for the epigenetic and Kinase libraries were collected for genomic DNA extraction, ensuring over 400x coverage of the libraries used for screening. Genomic DNA extraction was performed using a DNeasy Blood & Tissue Kit (Qiagen, Cat#69506) and quantified using a Nanodrop instrument. The sgRNA sequences were amplified via PCR using NEB Q5 polymerase (New England Biolabs, cat# M0491S). Purification of PCR products was done using AMPure XP SPRI beads (Beckman Coulter, Cat# A63881), and quantification was carried out using a Qubit dsDNA HS assay (Thermo Fisher Scientific, Cat# Q32851). Sequencing encompassed 4.5 million reads for transcription factor library and 5 million reads for the epigenetic and Kinase libraries on an Illumina HiSeq sequencer at the Hartwell Center Genome Sequencing Facility at St. Jude Children’s Research Hospital. NGS sequencing was carried out employing single-end, 100-cycle sequencing on a NovaSeq 6000 (Illumina). Validation to affirm gRNA presence and representation was conducted using calc_auc_v1.1.py (https://github.com/mhegde/) and count_spacers.py. CRIPSR data analysis was performed by using MAGeCK-VISPR software.

### RNA extraction from tumor tissues

Mice were humanely euthanized using CO2. Tumor samples were excised promptly and flash-frozen in liquid nitrogen, then preserved at −80°C until future utilization. A portion of the frozen sample was introduced into 500 µl RLT buffer (containing β-mercaptoethanol) and homogenized using a homogenizer (Pro scientific). The resulting mixture underwent centrifugation at 13000 rpm, and the supernatant was carefully transferred to a gDNA eliminator spin column, and further processed according to the manufacturer’s instructions for RNA extraction by utilizing the RNeasy Plus Mini Kit (Qiagen, ref. #74136).

### RNA-seq and analysis

Total RNA from cells and tumor tissues were performed using the RNeasy Mini Kit (Qiagen) according to the manufacturer’s instructions. Paired-end sequencing was performed using the High-Seq platform with 100bp read length. Total stranded RNA sequencing data were processed by the internal AutoMapper pipeline. Briefly the raw reads were firs trimmed (Trim-Galore version 0.60), mapped to human genome assembly (GRCh38) (STAR v2.7) and then the gene level values were quantified (RSEM v1.31) based on GENCODE annotation (v31). For mouse tumors, the raw reads were first trimmed (Trim Galore version 0.60), mapped to mouse genome assembly (GRCm38, mm10) (STAR v2.7) and then the gene level values were quantified (RSEM v1.31) based on GENCODE annotation (VM22). Low count genes were removed from analysis using a CPM cutoff corresponding to a count of 10 reads and only confidently annotated (level 1 and 2 gene annotation) and protein-coding genes are used for differential expression analysis. Normalization factors were generated using the TMM method, counts were then transformed using voom and transformed counts were analyzed using the lmFit and eBayes functions (R limma package version 3.42.2). Then Gene set enrichment analysis (GSEA) was carried out using gene-level log2 fold changes from differential expression results against gene sets in the Molecular Signatures Database (MSigDB 6.2) (gsea2 version 2.2.3). After mapping RNA-seq data, rMATS v4.1.0 was used for RNA alternative splicing analysis by using the mapped BAM files as input. Specifically, five different kinds of alternative splicing events were identified, i.e., skipped exon (SE), alternative 5’-splicing site (A5SS), alternative 3’-splicing site (A3SS), mutually exclusive exon (MXE) and intron retention (RI). To keep consistent, the same GTF annotation reference file for mapping was used for rMATS. For stranded RNA-seq data, the argument “--libType fr-firststrand” was applied. To process reads with variable lengths, the argument “--variable-read-length” was also used for rMATS. To select statistically significantly differential splicing events, the following thresholds were used: FDR <0.05 and the absolute value of IncLevelDifference > 0.1. For visualization, the IGV Genome Browser was used to show the sashimi plots of splicing events.

### Assay for Transposase-Accessible Chromatin using sequencing (ATAC-seq), ATAC-seq alignment, peak-calling and annotation

Library preparations for ATAC-seq were based on a published protocol with minor modifications ^100,101^. Briefly, freshly cultured SKNAS cells (100,000 per sample, naïve and two resistant ones) were harvested and washed with 150 μl ice-cold Dulbecco’s Phosphate-Buffered Saline (DPBS) containing protease inhibitor (PI). Nuclei were collected by centrifugation at 500 g for 10 minutes at 4 °C after cell pellets were resuspended in lysis buffer (10 mM Tris-Cl pH 7.4, 10 mM NaCl, and 3 mM MgCl_2_ containing 0.1% NP-40 and PI). Nuclei were incubated with Tn5 transposon enzyme in transposase reaction mix buffer (Illumina, cat# 20034197) for 30 min at 37 °C. DNAs were purified from the transposition sample by using Mini-Elute PCR purification kit (cat#28004, Qiagen) and measured by Qubit. Polymerase chain reaction (PCR) was performed to amplify with High-Fidelity 2X PCR Master Mix [72°C/5 mins + 98 °C /30 s + 12 × (98 °C /10 s + 63 °C/30 s + 72 °C/60 s) + 72 °C/5 min]. The libraries were purified using the Mini-Elute PCR purification kit (Qiagen). ATAC-seq libraries were pair-end sequenced on HiSeq4000 (Illumina) in the Hartwell Center at St Jude Children’s Research Hospital, Memphis, TN, USA.

The ATAC-seq raw reads were aligned to the human reference genome (hg38) using BWA ^102^ to and then marked duplicated reads with Picard (version 1.65), with only high-quality reads kept by samtools (version 1.3.1, parameter ‘‘-q 1 -F 1024’’)^103^. Reads mapping to mitochondrial DNA were excluded from the analysis. All mapped reads were offset by +4 bp for the + strand and -5 bp for the – strand ^100^. Peaks were called for each sample using MACS2 ^104^ with parameters “-q 0.01 –nomodel – extsize 200 –shift 100”. Peaks were merged for the same cell types using BEDtools^105^. Peak annotation was performed using HOMER^106^. All sequencing tracks were viewed using the Integrated Genomic Viewer (IGV 2.3.82) ^107^.

### CUT&Tag and analysis

CUT&Tag DNAs from SKNAS-naïve, IR1 and IR2 cells and PDX-SJNB14-ctrl and PDX-SJNB14-IR tumors were prepared by following the protocol as described previously^108^ (https://www.protocols.io/view/bench-top-cut-amp-tag-bcuhiwt6?step=1) with minor modifications. Briefly, for SJNB14-ctrl and SJNB14-IR tumor, nuclei extraction is following the protocol of isolate nuclei from frozen tissues (https://support.missionbio.com/hc/en-us/article_attachments/4421562098967). For SKNAS-naïve, IR1 and IR2 cells were washed with wash buffer (20 mM HEPES pH 7.5; 150 mM NaCl; 0.5 mM Spermidine; 1× Protease inhibitor cocktail). Nuclei were isolated with cold NE1 buffer (20 mM HEPES–KOH, pH 7.9; 10 mM KCl 0.1%; Triton X-100; 20% Glycerol, 0.5 mM Spermidine; 1x Protease Inhibitor) for 10 min on ice. Nuclei were collected by 600 x g centrifuge and resuspended in 1ml washing buffer containing with 10 µL of activated concanavalin A-coated beads (Bangs laboratories, BP531) at RT for 10 min. Bead-bound nuclei were collected with placing tube on magnet stand and removing clear liquid. The nuclei bound with bead were resuspended in 50 µL Dig-150 buffer (20 mM HEPES pH 7.5; 150 mM NaCl; 0.5 mM Spermidine; 1× Protease inhibitor cocktail; 0.05% Digitonin; 2 mM EDTA) and incubated with a 1:50 dilution of H3K27ac (Abcam, ab4729; RRID:AB_2118291) overnight at 4 °C. The unbound primary antibody was removed by placing the tube on the magnet stand and withdrawing the liquid. The primary antibody bound nuclei bead was mixed with Dig-150 buffer 100uL containing guinea pig anti-Rabbit IgG antibody (Antibodies, ABIN101961; RRID:AB_10775589) 1:100 dilution for 1 hour at RT. Beads bound nuclei were washed using the magnet stand 3× for 5 min in 1 mL Dig-150 buffer to remove unbound antibodies. A 1:100 dilution of pA-Tn5 adapter complex was prepared in Dig-300 buffer (20 mM HEPES, pH 7.5, 300 mM NaCl, 0.5 mM Spermidine, 0.05% Digitonin, 1× Protease inhibitor cocktail). After removing the liquid on the magnet stand, 100 µL mixture of pA-Tn5 and Dig-300 buffer was added to the nuclei bound beads with gentle vortex and incubated at RT for 1 h. After 3× 5 min in 1 mL Dig-300 buffer to remove unbound pA-Tn5 protein, nuclei were resuspended in 250 µL Tagmentation buffer (10 mM MgCl_2_ in Dig-300 buffer) and incubated at 37 °C for 1 h. EDTA 10 µL of 0.5 M, 3 µL of 10% SDS and 2.5 µL of 20 mg/mL Proteinase K were added to stop tagmentation and incubated at 55 °C for 1 hour. DNA libraries were then purified with SPRIselect beads (Beckman Coulter, B23318) following manufacture instruction and then dissolved in water for Illumina sequencing.

The sequencing raw reads were aligned to the human reference genome (hg38) using BWA (version 0.7.12; BWA aln+sampe for CUT&Tag data). Duplicate reads were marked and removed by Picard (version 1.65). For CUT&Tag, only properly paired uniquely mapped reads were extracted by samtools (version 1.3.1 parameters used were -q 1 -f 2 -F 1804) for calling peaks and generating bigwig file. Narrow peaks were called by MACS2 (version 2.2.7.1) with parameters of “ -t cut_tag_file -q 0.05 -f BED --keep-dup all”. Peak regions were defined to be the union of peak intervals in replicates from control or treated cells respectively. For peak overlap analysis, mergeBed (BEDtools version 2.25.0) was used to combine overlapping regions from multiple peak sets into a new region and then a custom script was used to summarize common or distinct peaks and visualize in a Venn diagram. Promoter regions was defined as the regions 1.0 kb upstream and 1.0 kb downstream of the transcription start sites based on the human RefSeq annotation (hg38). Genomic feature annotation of peaks was done by annotatePeaks.pl, a program from the HOMER suite (v4.8.3, http://homer.salk.edu/homer/). We used genomeCoverageBed (BEDtools 2.25.0) to produce genome-wide coverage in BEDGRAPH file and then converted it to a bigwig file by bedGraphToBigWig. The bigwig files were scaled to 15 million reads to allow comparison across samples. To show average of several replicates as a single track in the browser, the bigwig files were merged to a single average bigwig file using UCSC tools bigWigtoBedGraph, bigWigMerge and bedGraphToBigWig. The Integrated Genomics Viewer (IGV 2.3.82) was used for visual exploration of data. The HOMER software was used to perform de novo motif discovery as well as a check the enrichment of known motifs in a set of given peaks. Motif density histograms were created using HOMER for target regions. Background regions were generated by selecting DNA sequences of equal length at 10 kb downstream of the target regions. The motif density at target regions was normalized to that at the control regions.

### Flow cytometry for immune cell profiling of transgenic TH-MYC/ALK^F1178L^ tumors

To generate tumor single cell suspensions, tumors were excised, mechanically dissociated and transferred to basal medium containing 0.1% Collagenase Type IV and 150 mg/mL DNase I, incubated for 30-minute shaking at 37°C, then passed through a 70-μm nylon mesh. Cells were treated with ACK red blood cell lysis buffer and resuspended in PBS prior to further analysis. Cells were treated with a 1:100 dilution of CD16/32 blocking antibody (clone 2.4G2, Tonbo) for 10 min at 4°C, then stained for 30 min at 4°C with the following antibodies/dyes: B220 AF647 (clone RA3-6B2, BioLegend, lot# B301694, dilution 1:200) for B cells, CD3 PE/Dazzle594 (clone 17A2, BioLegend, lot# B307674, dilution 1:200) for T cells, CD4 BV570 (clone RM4-5, BioLegend, lot# B341697, dilution 1:200) for CD4 T cells, CD8a BV711 (clone 53-6.7, BioLegend, lot# B100747, dilution 1:200) for CD8 T cells, CD45 PE/Cy5 (clone 30-F11, BioLegend, lot# B340539, dilution 1:200), CD49b BV605 (clone HMα2S, BD Biosciences, lot# 1305998, dilution 1:200) for NK cells, Ghost Dye Violet 510 viability dye (Tonbo, 1:400). Data were acquired with a Cytek Aurora spectral flow cytometer and analyzed in FlowJo v10 (Treestar).

### Flow cytometry for NK cells in c-MYC tumors implanted in Rag2^-/-^ mice

Tumors were excised, mechanically dissociated and transferred to basal medium containing 0.1% Collagenase Type IV (Worthington Biochemical Corporation, Cat#CLS-4) and 150 mg/mL DNase I (Promega, Cat# M6101), incubated for 30-minute shaking at 37°C, then passed through a 70-μm nylon mesh. Cells were treated with ACK red blood cell lysis buffer and resuspended in PBS prior to further analysis. Cells were incubated in C-RPMI containing Cell Stimulation Cocktail with protein transport inhibitors (eBioscience, cat# 00-4975-93) or protein transport inhibitors alone (eBioscience, cat# 00-4980-93) for 4hr at 37°C. Cells were then treated with a 1:100 dilution of Fc receptor blocking antibody (clone 2.4G2, Tonbo) for 10 min at 4°C, and stained for 30 min at 4°C with the following antibodies/dyes: CD3 PE/Dazzle594 (clone 17A2, BioLegend, lot# B307674, dilution 1:200), CD45 PE/Cy5 (clone 30-F11, BioLegend, lot# B340539, dilution 1:200), CD49b BV605 (clone HMα2S, BD Biosciences, lot# 1305998, dilution 1:200), Ghost Dye Violet 510 viability dye (Tonbo, lot# D0870071921133, 1:400). Cells were fixed and permeabilized with Cyto-Fast Fix/Perm Buffer Set (BioLegend, cat# 426803) according to manufacturer’s instructions and stained with IFN-γ BV785 (clone XMG1.2, BioLegend, lot# B334622, dilution 1:100) for 30 min at 4°C. After washing cells, data were acquired with a Cytek Aurora spectral flow cytometer. and analyzed in FlowJo v10 (Treestar).

### In vivo plasma pharmacokinetics study

The plasma pharmacokinetic (PK) profile of the RBM39 degrader indisulam (E7070) was evaluated in normal female CB17/SCID mice (Taconic Biosciences), approximately 12 weeks in age. Indisulam (MedKoo, CAT# 201540, LOT# KB60825) was dissolved in 5% DMSO, 4.75% Tween 80 in saline, at 5 mg/mL for a 25 mg/kg free base equivalent dose as a 5 mL/kg IV bolus dose via the tail vein. Two survival blood samples were obtained from each mouse via retro-orbital plexus using Drummond EDTA 75mm capillary tubes, and a third final sample by cardiac puncture, all using KEDTA as the anticoagulant. Samples were obtained at various times up to 36 hours post-dose, immediately processed to plasma, and stored at -80 °C until analysis. Remaining dosing solution was submitted for verification of potency, and chemical and physical stability during the study period. Plasma samples were analyzed for indisulam using a qualified liquid chromatography – tandem mass spectrometry (LC-MS/MS) assay. Plasma calibrators and quality controls were spiked with solutions, corrected for salt content and purity as necessary, prepared in DMSO.

Plasma samples, 25 µL each, were protein precipitated with 100 µL of 30 ng/mL U-104 (MedChemExpress, CAT# HY-13513/CS-4495, LOT# 17909) in methanol as an internal standard (IS). A 3 µL aliquot of the extracted supernatant was injected onto a Shimadzu LC-20ADXR high performance liquid chromatography system via a LEAP CTC PAL autosampler. The LC separation was performed using a Phenomenex Kinetex EVO C18 (2.6 μm, 50 mm x 2.1 mm) column maintained at 50°C with gradient elution at a flow rate of 0.6 mL/min. The binary mobile phase consisted of UP water-methanol-formic acid, (90:10:0.1 v/v) and methanol-formic acid (100:0.1 v/v) in reservoir B. The initial mobile phase consisted of 20% B with a linear increase to 50% B in 1.5 min followed by a 1.0-minute hold at 50% B. The column was then rinsed for 2.0 min at 100% B and then equilibrated at the initial conditions for 2.0 min for a total run time of 6.5 min. Under these conditions, the analyte and IS eluted at 1.63 and 1.07 min, respectively. Analyte and IS were detected with tandem mass spectrometry using a SCIEX QTRAP 5500 in the negative ESI mode and the following mass transitions were monitored: indisulam 384.0 → 320.2, U-104308.1 → 171.1. The method qualification and bioanalytical runs all passed acceptance criteria for non-GLP assay performance. A linear model (1/X2 weighting) fit the calibrators across the 0.250 to 500 ng/mL range, with a correlation coefficient (R) of ≥ 0.9993. The lower limit of quantitation (LLOQ), defined as a peak area signal-to-noise ratio of 5 or greater verses a matrix blank with IS, was 0.250 ng/mL. Sample dilution integrity was confirmed, and no matrix effects were observed in blank experimental CB17/SCID plasma. The intra-run precision and accuracy was ≤ 6.32% CV and 94.7% to 105%, respectively.

Plasma concentration-time (Ct) data for indisulam were grouped by matrix and nominal time point and summary statistics calculated. The arithmetic mean Ct values were subjected to noncompartmental analysis (NCA) using Phoenix WinNonlin 8.1 (Certara USA, Inc., Princeton, NJ). The IV bolus model was applied, and area under the Ct curve (AUC) values were estimated using the “linear up log down” method. The terminal phase was defined as at least three time points at the end of the Ct profile, and the elimination rate constant (Kel) was estimated using an unweighted log-linear regression of the terminal phase. The terminal elimination half-life (T1/2) was estimated as 0.693/Kel, and the AUC from time 0 to infinity (AUCinf) was estimated as the AUC to the last time point (AUClast) + Clast (predicted)/Kel. Other parameters estimated included observed maximum concentration (Cmax), time of Cmax (Tmax), concentration at the last observed time point (Clast), time of Clast (Tlast), back extrapolated initial concentration (C0), clearance (CL = Dose/AUCinf), and volume of distribution at steady state (Vss). A clinically relevant dose (CRD) for mice was estimated from unbound plasma PK and exposure. The CRD was defined as the mouse dose regimen achieving a predicted mean unbound plasma average concentration over time (Cavg,t) similar to humans with the selected dose level and regimen of interest. The Cavg,t was calculated as the AUC from 0 to time t divided by time t. Dose proportional, linear, and time-invariant PK for mice was assumed. Human and mouse plasma protein binding were assumed to be similar since data were not available. This corresponds to the clinical relevance approach proposed by Spilker^109^ which uses unbound plasma average steady state concentrations to define CRDs. Some latitude in dose rounding was permitted in the CRD recommendation, and an unbound exposure within 2-fold of the clinical target was considered acceptable. Additional considerations influenced the final recommended mouse dose, including mouse dosing regimens prevalent in the literature and the tolerability of the compound in mice.

### Animals and Therapies

Sex as a biological variable. Both genders of Th-MYCN/ALK^F1178L^, C57BL/6, Rag2^-/-^ and NSG mice were used. All murine experiments were done in accordance with a protocol approved by the Institutional Animal Care and Use Committee of St. Jude Children’s Research Hospital. Subcutaneous xenografts were established in 4-6 weeks old CB17 SCID mice (CB17 scid, Taconic) or NOD.Cg-Prkdcscid Il2rgtm1Wjl/SzJ (NOD scid gamma (NSG, St Jude Children’s Research Hospital, bred in house ARC) mice by implanting 5 x10^6^ cells (SK-N-AS and SIMA) in Matrigel. Patient derived xenograft (SJNB14 PDX) and mouse derived xenograft (C-MYC MDX) tumors were sectioned into small fragments using a Razor blade (Electron Microscopy Sciences, Cat#71980), loaded into a syringe (21-gauge needle, BD: Ref:305167), and subcutaneously injected into mice. Following genotyping, *TH-MYCN*/*ALK*^F1178L^ mice were gender-segregated and assigned to imaging groups. Inclusion criteria were the presence of the *TH-MYCN* allele, heterozygosity for one *ALK*^F1178L^ allele, and an age range from 1 to 3 weeks postweaning. Either gender was used and noted. Criteria for analysis were the presence or absence of an ultrasound-detectable tumor (quantified by staff not directly involved in this study) at any point within a 7-week period, where mice were imaged once per week. To mitigate gender bias, both genders were employed in NSG, Rag2^-/-^, or transgenic TH-MYCN/ALK^F1178L^ mouse models. Tumor measurements were done weekly using electronic calipers for xenograft models, and volumes calculated as π/6 × d^3^, where d is the mean of two diameters taken at right angles. The starting treatment points and dosing schedules were documented in each figure legend. Indisulam was administered to CB17 SCID or NSG mice via tail vein injection, while TH-MYCN/ALK^F1178L^, C57BL/6, or Rag2^-/-^ mice received treatment through IP injection.

For ATR inhibitor gartisertib in combination with indisulam, once the tumor volume reached around 200 mm^3^, mice were randomly assigned to 4 groups (n=5 mice per group). Mice were administered with vehicle (15% Captisol /0.1M hydrochloride acid, oral gavage), indisulam (10 mg/kg, tail vein injection, 5 days on and 2 days off, for three weeks), gartisertib (10 mg/kg/day, oral gavage, once per week for three weeks), and indisulam (10 mg/kg, tail vein injection, for three weeks) combined with gartisertib (10 mg/kg/day, oral gavage once per week for three weeks) except the mice that reached humane endpoints. The groups with indisulam alone and combination therapy were given another two-week cycle treatment when the tumors in mice with indisulam alone treatment relapsed. The tumor volume and mice weight were measured twice in a week. Mice were euthanized when the tumor volume reached 20% of the body weight or the mice became moribund.

### Ultrasound imaging of transgenic TH-MYCN/ALK^F1178L^ mouse

Fur was removed from the ventral side of each animal using Nair. Technicians in the St. Jude Center for In Vivo Imaging and Therapeutics performed ultrasound scanning on mice weekly using VEVO-3100 and determined tumor volumes using VevoLAB 5.7.1 software. All ultrasound data were acquired in a blinded fashion.

### Quantification and statistical analysis

All data in this study are displayed as the mean ± SEM, unless indicated elsewise. Comparison between two groups was determined using Student’s t-test. Wilcoxon rank sum test (two-sided) was used to compare the tumor volumes between two groups at every time point. P-values across multiple time points were adjusted for multiple comparison using the Benjamini-Hochberg method. Kaplan-Meier survival was analyzed using log-rank (Mantel-Cox) method in Prism program.

## Resource availability

### Lead contact

Further information and requests for resources and reagents should be directed to the lead contact, Jun Yang.

### Materials availability

All CRISPR libraries, plasmids, mouse allograft models and cell lines generated in this study are available from the lead contact Dr. Jun Yang with a completed material transfer agreement.

### Data and code availability

The RNA-seq data, ATAC-seq data, CUT&Tag data have been deposited in GEO database.

(1) The GSE251920 SuperSeries is composed of the following SubSeries:

GSE251915 [CUT&Tag of H3K27Ac for SKNAS and SJNB14 models]

GSE251918 [RNA-seq for TH-MYCN/ALK^F1178L^ and SJNB14 models]

The reviewer access

link: https://www.ncbi.nlm.nih.gov/geo/query/acc.cgi?&acc=GSE251920&token=sxopsaqanvahlqn

(2) GSE164505 [RNA-seq for SIMA and SKNAS models, ATAC-seq for SKNAS model] Link: https://0-www-ncbi-nlm-nih-gov.brum.beds.ac.uk/geo/query/acc.cgi?acc=GSE164505

## ACKNOWLEDGEMENTS

We thank the staff in St. Jude Hartwell Center for their dedication and expertise. We thank Dr. Dongli Hu for his monthly mycoplasma testing and STR assay. This work was partly supported by American Cancer Society-Research Scholar (130421-RSG-17-071-01-TBG, J.Y.) and National Cancer Institute (1R01CA229739-01, 1R01CA266600-01A1, J.Y.), Comprehensive Cancer Center core grant P30 CA021765, and the American Lebanese Syrian Associated Charities (ALSAC). The content is solely the responsibility of the authors and does not necessarily represent the official views of the National Institutes of Health. The authors have declared that no conflict of interest exists.

## AUTHOR CONTRIBUTIONS

S.S. performed animal experiments and CRISPR screening. J.F performed ATAC-seq and CUT&TAG. H.J. performed bioinformatics analysis. L.V and P.T provided transgenic TH-MYCN/ALK^F1178L^ mice, immune cell infiltration analysis. Q.W. analyzed the GD2 expression with help from L.H. and performed quality control of indisulam. W.Q. examined RBM39 mutations. A.C. performed in vitro analysis of SK-N-AS response to indisulam. C.L.M provided PDX. C.L.M, M.A.W with W.V.C and B.B.F III. performed PK study. J.P.C, J.A.S and S.M.P-M provided CRISPR library expansion, and pooled screen analysis. R.T., G.T., T.C. and M.J performed small animal imaging and analysis. T. L provided anti-GD mAb., R. W generated Dbh-iCre/CAG-MYC allograft model. P.T, A.M.D, and J.Y. conceived the project. J.Y. designed the study, analyzed data, and wrote the manuscript with help from all authors.

