## Supplemental Figures for "RBM39 degrader invigorates natural killer cells to eradicate neuroblastoma despite cancer cell plasticity"

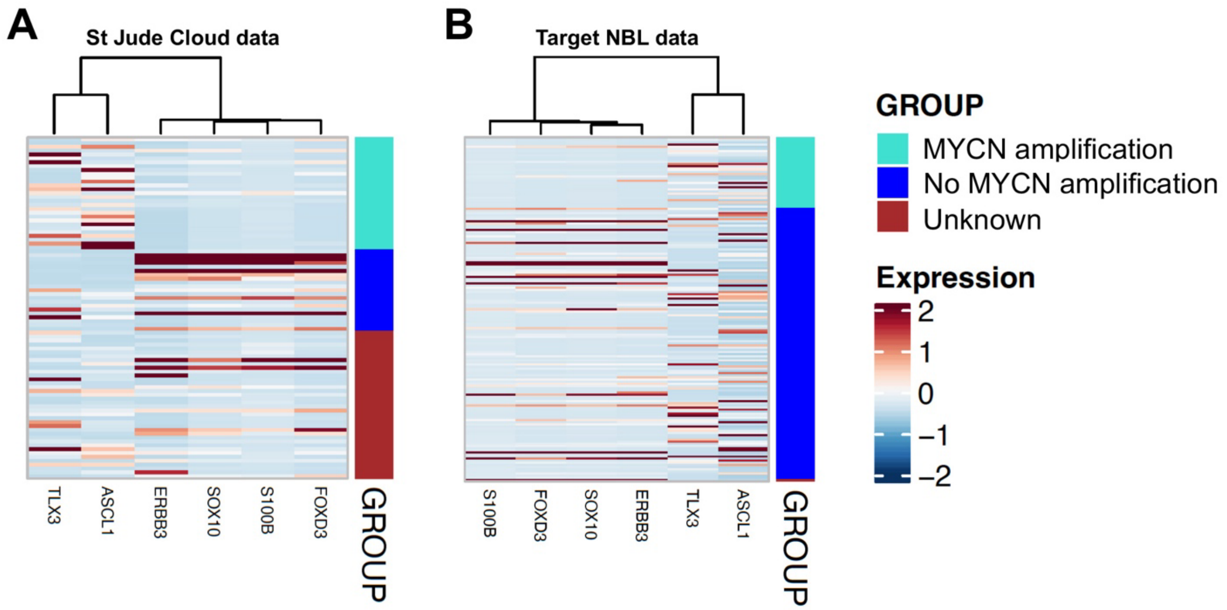


**Supplemental Figure S1. Expression of Schwann cell precursor in human neuroblastomas.**

Raw RNA-seq data requested from Pediatric Cancer Genome Project (PCGP) (A) and NCI's Therapeutically Applicable Research To Generate Effective Treatments (TARGET, https://ocg.cancer.gov/programs/target) (B) were processed by internal AutoMapper pipeline (described in method section of RNA-seq and analysis). TPM (transcript per million) matrix of neuroblastoma samples from both studies were extract respectively to generate heatmaps for SCP signature. MYCN amplification status were annotated in color bars. Heatmap pseudocolor indicated z-score of log2TPM.

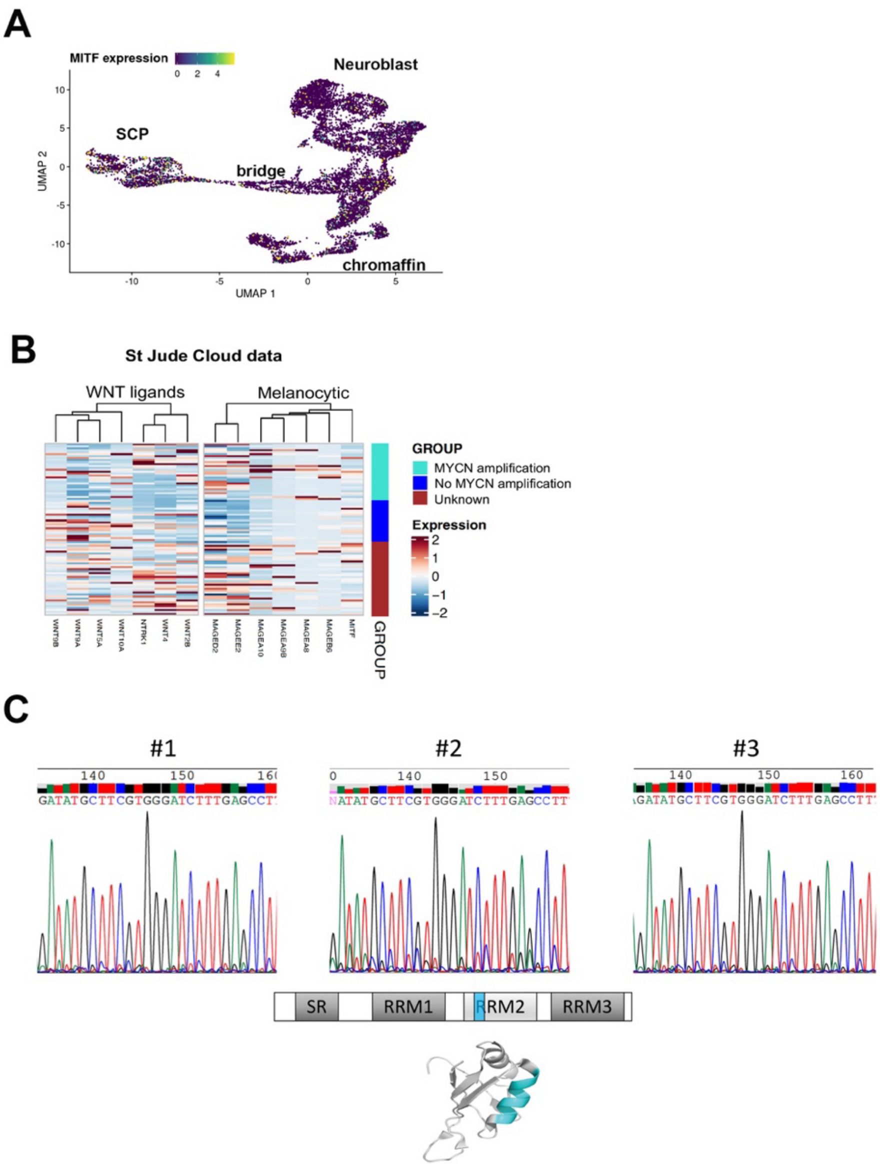


**Supplemental Figure S2. Deconvolution of gene signatures in indisulam-resistant SJNB14 PDX.** (A) Low *MITF* expression in the SCP cell population, bridge cells, neuroblasts and Chromaffin cells from study (https://adrenal.kitz-heidelberg.de/developmental_programs_NB_viz/). (B) Raw RNA-seq data requested from Pediatric Cancer Genome Project (PCGP) were processed by internal AutoMapper pipeline (described in method section of RNA-seq and analysis). TPM matrix of neuroblastoma samples were extract respectively to generate heatmaps for WNT ligands and melanocytic signature. MYCN amplification status were annotated in color bars. Heatmap pseudocolor indicated z-score of log2TPM. (C) Sanger DNA sequencing verifies that there are no mutations at the hotspot sequence of RBM39 in three resistant tumors. The hotspot was indicated by the blue color in RRM2 motif of RBM39.

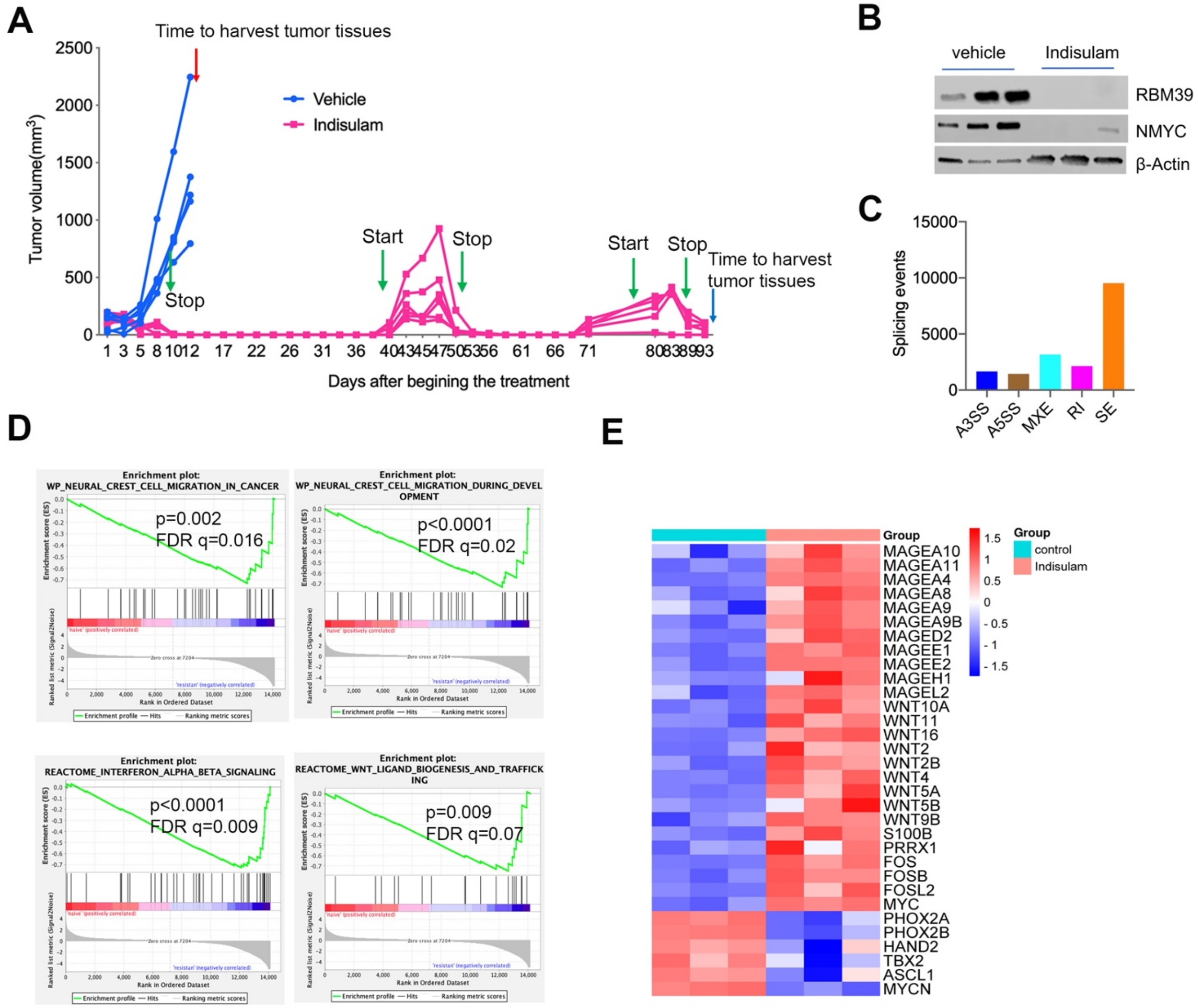


**Supplemental Figure S3. SIMA tumor cells undergo cell state alterations during indisulam therapy.** (A) Tumor growth curve for SIMA implanted in NSG mice undergoing repeated cycles (2-week as one cycle) of treatment with 25mg/kg indisulam, 5 days on, two ways off. (B) Western blot analysis of SIMA xenograft tissues treated with indisulam during the third cycle of treatment, with indicated antibodies. (C) Alternative splicing events induced by indisulam during third cycle of treatment. (D) GSEA shows the gene signatures significantly upregulated in resistant tumors vs naïve tumors. (E) Heatmap showing that the expression changes of melanocytic markers, Wnt ligands, MES and ADRN transcriptional factors in naïve vs resistant tumors.

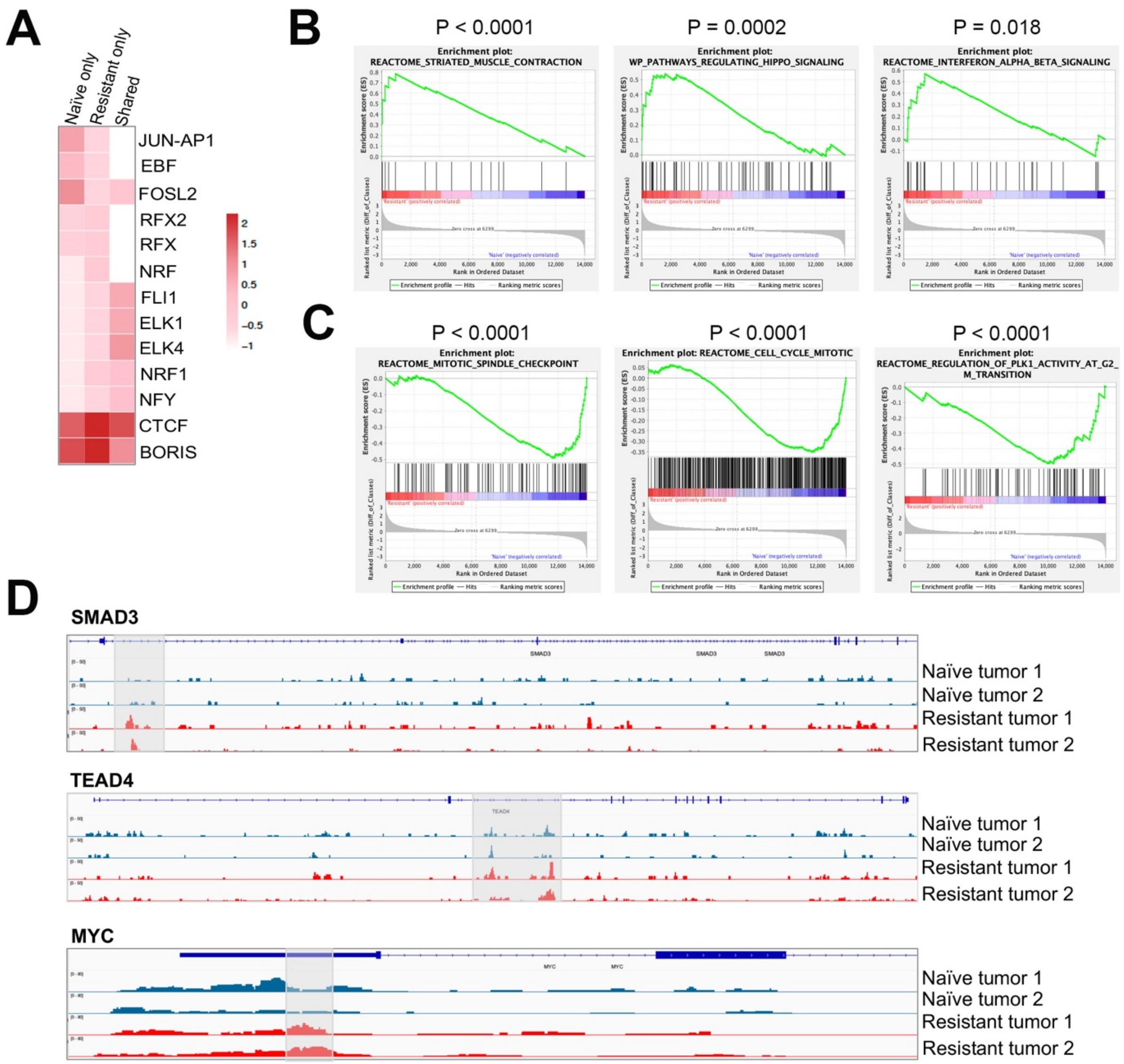


**Supplemental Figure S4. Epigenetic reprogramming of MYCN–amplified SJNB14 tumor cells after they developed full resistance to indisulam therapy after repeated treatments.** (A) Heatmap indicates the binding motifs for transcriptional factors enriched in naïve, resistant and both based on Homer Motif analysis of H3K27Ac peaks. (B, C) GSEA results for the H3K27Ac upregulated (B) and downregulated (C) at the genes in resistant tumors. (D) Snapshots by IGV showing the H3K27Ac peaks (highlighted in grey color) upregulated at the genomic loci of MES CRC TFs in resistant tumors.

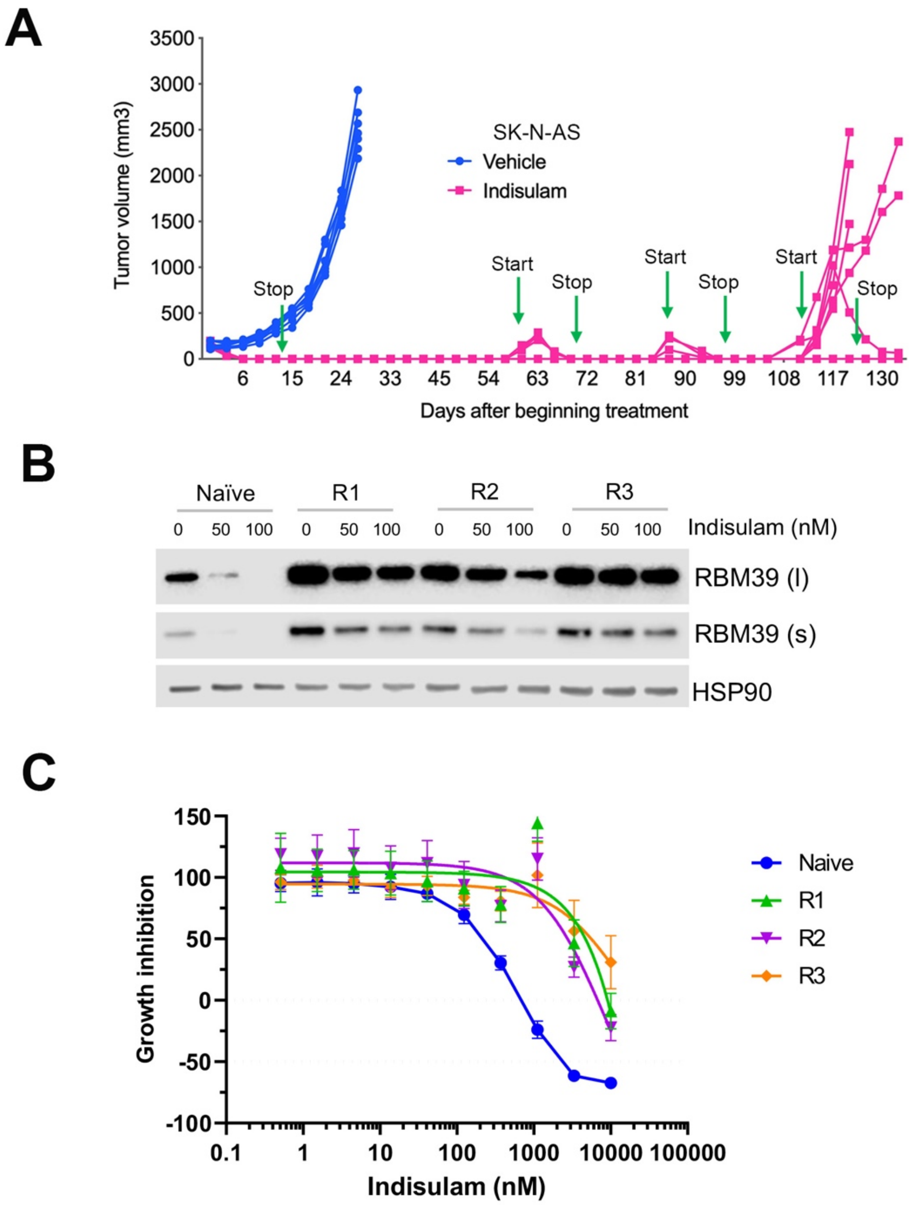


**Supplemental Figure S5. SK-N-AS tumor cells develop full resistance to indisulam therapy after repeated treatments.** (A) Tumor growth curve for SK-N-AS implanted in NSG mice undergoing repeated cycles (2-week as one cycle) of treatment with 25mg/kg indisulam, 5 days on, two ways off. (B) Western blot analysis of SK-N-AS naïve and indisulam-resistant cells treated with different concentrations of indisulam with indicated antibodies. L, long exposure of the gel; s, short exposure of the gel. (C) Growth inhibition curve of SK-N-AS naïve and indisulam-resistant cells assessed by Prestoblue assay.

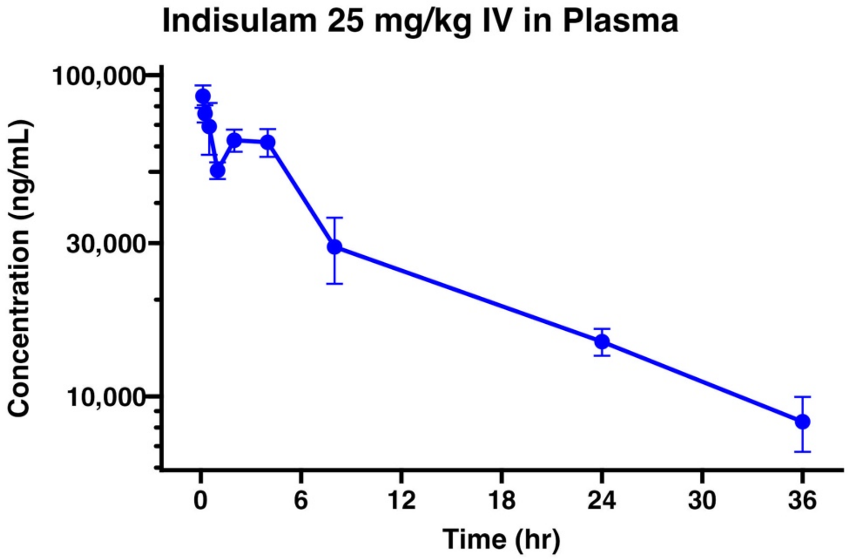


**Supplemental Figure S6. Plasma concentration-time profile of indisulam from pharmacokinetic study.** Indisulam concentrations in plasma of CB17/SCIG mice were assessed by mass spectrometry over time after a 25 mg/kg bolus tail vein injection of indisulam. Data points are shown as mean and standard deviation.

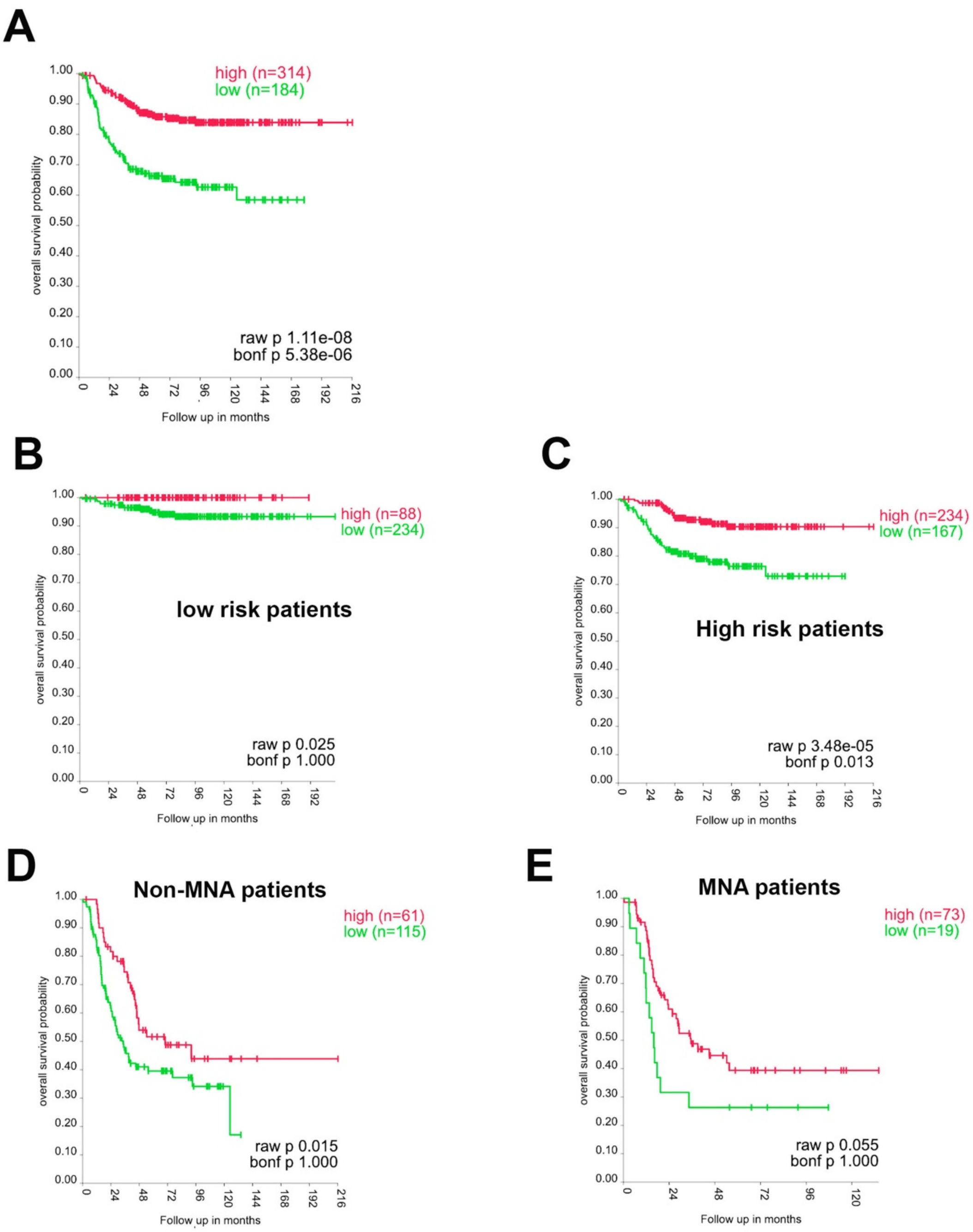


**Supplemental Figure S7. High expression of *NCR1* is correlated with a better survival of neuroblastoma patients.** (A) Kaplan-Meier overall survival for neuroblastoma patients with high and low expression of *NCR1* gene in SEQC cohort (GSE62564). (B, C) Kaplan-Meier overall survival for low risk (B) and high risk (C) neuroblastoma patients with high and low expression of *NCR1* gene in SEQC cohort (GSE62564). (D, E) Kaplan-Meier overall survival for non-MYCN amplified (D) and MYCN amplified (E) neuroblastoma patients with high and low expression of *NCR1* gene in SEQC cohort (GSE62564).

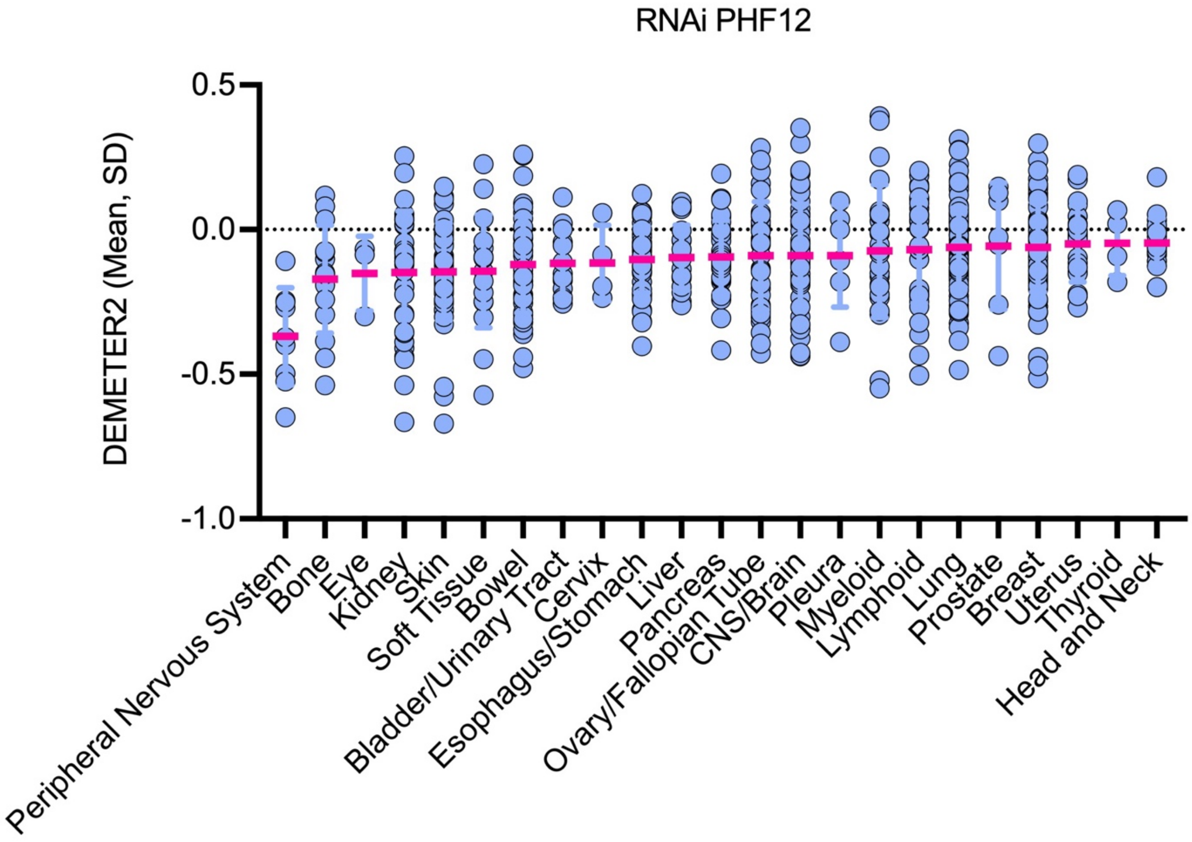


**Supplemental Figure S8. PHF12 is a neuroblastoma dependency gene.** The DEMETER2 score of an RNAi screen across 25 cancer lineages. A lower DEMETER2 score indicates a higher likelihood that a gene of interest is essential in a given cell line. The DETERM2 score was represented by mean ±SD. Data was downloaded from Depmap.org and plotted by using PRISM.
